# Cell-type specific requirement for pyruvate dehydrogenase in hematopoiesis and leukemia development

**DOI:** 10.1101/2021.05.04.442528

**Authors:** Sojeong Jun, Swetha Mahesula, Thomas P. Mathews, Misty S. Martin-Sandoval, Zhiyu Zhao, Elena Piskounova, Michalis Agathocleous

**Author notes:** Sandra and Edward Meyer Cancer Center and Department of Dermatology, Weill Cornell Medicine, New York, New York (Elena Piskounova).

## Abstract

Cancer cells have different metabolic requirements as compared to their corresponding normal tissues. This is thought to reflect metabolic reprogramming during transformation. An alternative possibility is that some metabolic requirements of cancer cells reflect a maintenance of the metabolism of the specific normal cell type from which cancer cells originate. Here, we investigate this hypothesis by comparing glucose use in normal hematopoiesis and in leukemia. T cell progenitors in the thymus were glucose avid and oxidized more glucose in the tricarboxylic acid (TCA) cycle through pyruvate dehydrogenase (PDH) as compared to hematopoietic stem cells (HSCs) or other hematopoietic cells. PDH deletion reduced the number of double positive (DP) T cell progenitors but had no effect on HSCs, myeloid progenitors and other hematopoietic cells we examined. PDH deletion blocked the development of T cell leukemia from *Pten-*deficient DP progenitors, but not the development of a myeloid neoplasm from *Pten-*deficient HSCs or myeloid progenitors. Therefore, the requirement of glucose oxidation for leukemia development is inherited from the normal cell of origin and occurs independently of the driver genetic lesion. PDH was not required *in vivo* to generate acetyl-CoA or maintain levels of TCA cycle metabolites but to prevent pyruvate accumulation and to maintain glutathione levels and redox homeostasis.

## INTRODUCTION

Cancer cell metabolism is shaped both by oncogenic processes and by the metabolism of the normal tissue of origin (Vander Heiden and DeBerardinis, 2017). Previous studies found that the metabolism of cancer cells is more similar to the metabolism of the corresponding normal tissue than to cancer cells from other tissues (Hu et al., 2013; Mayers et al., 2016; Yuneva et al., 2012), and that cancer cell metabolism is regulated by lineage-specific transcription factors (Huang et al., 2018). Yet there is also clear evidence for cancer-specific metabolism from studies showing widespread metabolic changes in cancers as compared to the normal tissue that the cancer originates from (Vander Heiden and DeBerardinis, 2017). Given that normal tissues consist of different cell types, and that cancer cells usually originate from one or a small subset of cell types within a tissue, it is possible that even metabolic features that are different between cancers and normal tissue are simply inherited from the normal cell type that originated the cancer. In this scenario, cancer driver mutations maintain, rather than reprogram, the metabolism of the normal cell which originates the cancer. Parallels between normal stem or progenitor cell and cancer cell metabolism suggest this might be the case. For example, loss of the mitochondrial pyruvate carrier MPC1 promotes intestinal stem cell proliferation (Schell et al., 2017) and colorectal cancer initiation (Bensard et al., 2020). If a metabolic requirement for cancer development is inherited from the cell of origin, the prediction is that it should be present in cancer cells and their normal cells of origin, but not in cancers of the same tissue type, driven by the same genetic lesion but which originate from a different stem or progenitor cell type. To test this idea, we investigated glucose metabolism in hematopoietic cell types and in their corresponding leukemia cells.

Normal stem or progenitor cells are thought to rely on glycolytic metabolism in several tissues (Agathocleous et al., 2012; Flores et al., 2017; Zheng et al., 2016). Previous work suggested that glucose uptake and glycolysis are high in hematopoietic stem cells (HSCs) and decline with differentiation (Cabezas-Wallscheid et al., 2014; Guo et al., 2018; Miharada et al., 2011; Simsek et al., 2010; Takubo et al., 2013; Wang et al., 2014). HSCs may have low mitochondrial activity, reside in a hypoxic niche (Takubo et al., 2010) and rely on glycolysis rather than on glucose oxidation (Simsek et al., 2010; Takubo et al., 2013; Vannini et al., 2016). Inhibition of Pdk1/2/4, which increases glucose oxidation by activating pyruvate dehydrogenase (PDH), impairs HSC reconstitution ability after transplantation (Halvarsson et al., 2017; Takubo et al., 2013). Inhibition of PPAR-γ, which increases glycolysis, boosts HSC maintenance *in vitro* (Guo et al., 2018). Deletion of lactate dehydrogenase A (*Ldha*) impairs HSC reconstituting activity after serial transplantation (Wang et al., 2014) and inhibition of Ppm1k (Liu et al., 2018) and Meis1 (Kocabas et al., 2012) impair both HSC reconstituting activity and glycolysis. However other work reached different conclusions. Oxygen levels in HSC niches are similar or higher than those for restricted progenitors or other hematopoietic cells (Nombela-Arrieta et al., 2013; Spencer et al., 2014), and HSCs are not regulated by hypoxia-inducible factor (HIF) proteins (Guitart et al., 2013; Vukovic et al., 2016). HSCs have high mitochondrial mass relative to restricted progenitors (de Almeida et al., 2017). Deletion of components of the TCA cycle or electron transport chain, such as complex III (Anso et al., 2017), succinate dehydrogenase (Bejarano-Garcia et al., 2016) and fumarate hydratase (Guitart et al., 2017) impairs HSC activity by affecting survival, proliferation or differentiation. In *Foxo3-*deficient mice, impairment in oxidative metabolism and activation of glycolysis is associated with impaired HSC reconstituting activity (Rimmele et al., 2015). Proliferating HSCs may increase glucose consumption, link glycolysis with mitochondrial metabolism and increase oxidative phosphorylation as compared to quiescent HSCs (Chandel et al., 2016; Liang et al., 2020; Maryanovich et al., 2015; Takubo et al., 2013). These divergent conclusions raise the question of whether glucose feeds the TCA cycle in quiescent or in proliferating HSCs and different types of restricted hematopoietic progenitors. Understanding the role of glycolysis and glucose oxidation in hematopoiesis has been complicated by the fact that previous work assessed HSC and progenitor metabolism *in vitro* and did not directly test the idea that proliferating HSCs rely on glucose oxidation *in vivo*.

## RESULTS

### The thymus is glucose avid but HSCs and other hematopoietic cells are not

To assess glucose uptake *in vivo*, we injected mice with 2-NBDG, a fluorescent glucose analogue used to measure glucose uptake *in vivo* and *in vitro* (Yamada et al., 2007). HSCs (CD150^+^CD48^-^Lineage^-^Sca-1^+^c-Kit^+^ cells) and multipotent progenitors (MPPs, CD150^-^CD48^-^ Lineage^-^Sca-1^+^c-Kit^+^) showed one of the lowest 2-NBDG signal of all cell types we measured (**Figure 1a**), suggesting that they take up less glucose than most other hematopoietic cells. Among bone marrow hematopoietic cells, CD8^+^ T cells had the highest glucose uptake (**Figure 1a**), consistent with previous reports that T cells use high amounts of glucose *in vitro* and *in vivo* (Ma et al., 2019; Macintyre et al., 2014). The thymus, where T cell progenitors develop, was glucose avid as compared to the bone marrow and spleen **(Figure 1b)**, consistent with observations of high ^18^F-fluorodeoxyglucose uptake in the thymus in children (Ferdinand et al., 2004; Jerushalmi et al., 2009), who have more active thymopoiesis than adults. To measure glucose utilization with an alternative method, we infused mice with uniformly labeled ^13^C (U^13^C) glucose and measured its contribution to metabolites at steady-state 3 hours later (Marin-Valencia et al., 2012) **(Figure 1c)**. Glucose-derived carbons contributed to glycolysis and the TCA cycle at higher rates in the thymus than in the bone marrow *in vivo* **(Figure 1d)**, consistent with higher glucose uptake and utilization in T cell progenitors than in other hematopoietic progenitors.

**Figure 1.**
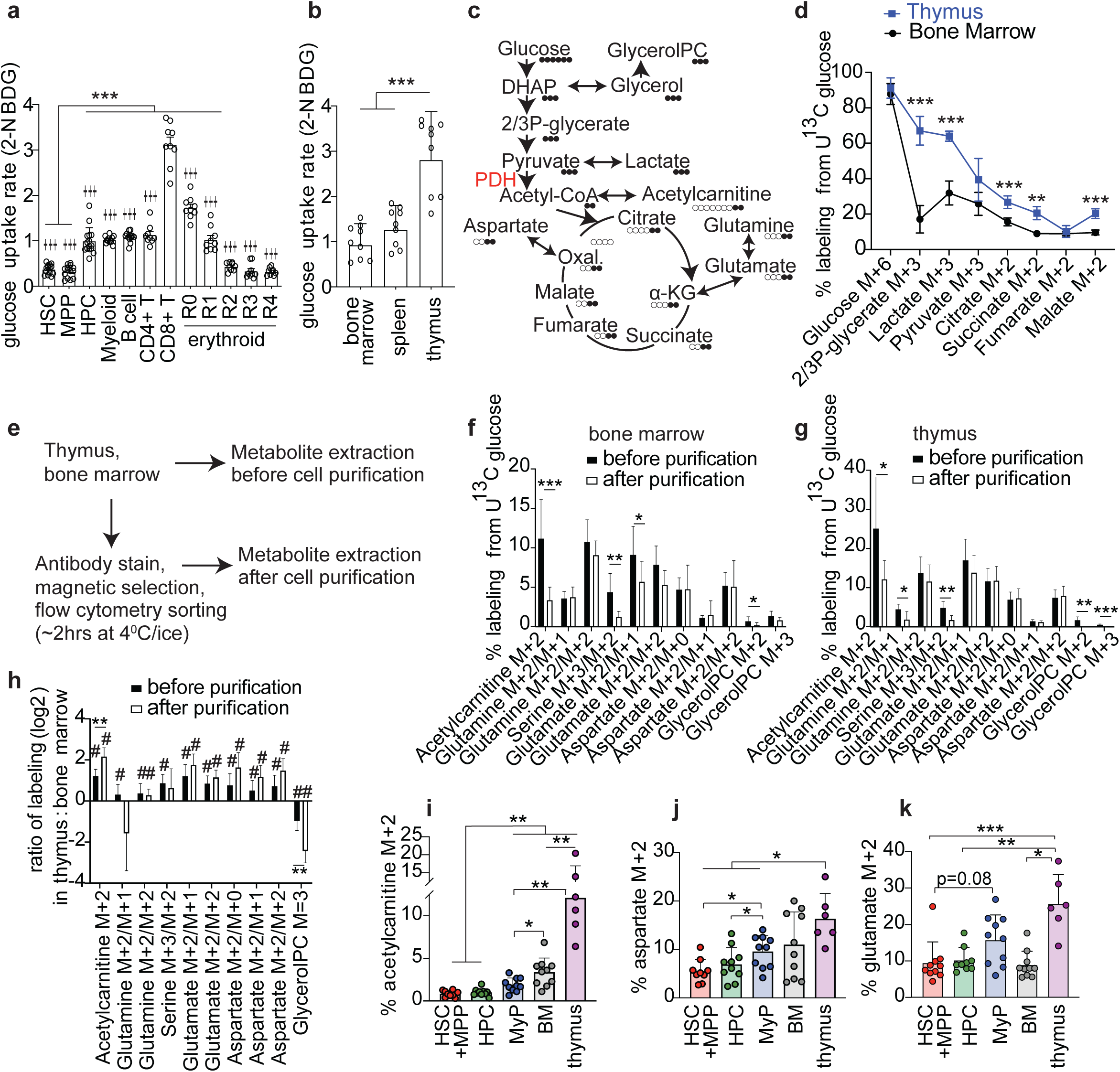
The thymus is glucose avid compared to HSCs and restricted hematopoietic progenitors in the bone marrow. **(a-b)** 2-NBDG median fluorescence intensity in different hematopoietic cell types in the bone marrow *in vivo* (n= 9-15 mice from 5 independent experiments, * Comparisons to HSCs, † comparisons to CD8^+^ T cells) or in whole bone marrow, spleen and thymus (n= 9 mice from 5 independent experiments). **(c)** Schematic of *in vivo* U^13^C-glucose infusion experiment depicting metabolites labeling patterns from glycolysis and the TCA cycle. **(d)** Fractional enrichment of metabolites in the mouse thymus and bone marrow after U^13^C-glucose infusion, normalized to % plasma glucose M+6 (n= 5 mice from 5 independent experiments). **(e-h)** Testing of the effect of cell purification on fractional enrichment after *in vivo* U^13^C-glucose infusion (n= 6-10 mice from 3 independent experiments). **(f-g)** Fractional enrichment for the indicated isotopomers in bone marrow or thymus cells before and after purification. The notation M+2/M+1 denotes labeling with 2 heavy carbons in the full molecule and 1 heavy carbon in the fragment used in the MS/MS transition. **(h)** Differences in isotopomer fractional enrichment in thymus *vs* bone marrow cells before and after cell purification. # denotes isotopomer fractional enrichment which was significantly different between thymus and bone marrow. * denotes isotopomer fractional enrichment whose relative levels between thymus and bone marrow changed after purification. **(i-k)** Enrichment of acetylcarnitine M+2, aspartate M+2, and glutamate M+2 in HSC+MPPs, HPCs, MyPs, bone marrow, and thymus cells purified from tissues directly after U^13^C-glucose infusion *in vivo*. 10,000 cells HSCs+MPPs or HPCs or MyPs, and 20,000-50,000 CD45+ bone marrow or thymus cells were used (n= 6-10 mice from 3 independent experiments). Each data point in all figures represents data from 1 mouse. All data represent mean ± s.d., or for (h) geometric mean ± geometric s.d. *p < 0.05, **p < 0.01, ***p < 0.001. Statistical significance was assessed with one-way ANOVA with Brown-Forsythe correction (a), one-way ANOVA (b), multiple t-tests (d), multiple t-tests, with the Welch correction for metabolites with unequal variance (f, g, h), and paired mixed-effects model (i-k). Multiple comparisons correction was performed by controlling the false discovery rate at 5% or by using multiple comparisons tests as described in the methods.

Understanding the metabolism of rare cell types in the body, such as tissue specific stem cells, has been limited by our inability to trace metabolites in small numbers of cells purified from tissues. Stable isotope tracing methods typically use cultured cells or bulk tissues (Jang et al., 2018). Methods for cells purified from tissues have been recently developed, including for ^13^C-glucose-derived labeling in 1-2 million T cells purified from spleen (Ma et al., 2019) and to measure labeling in amino acids derived from hydrolyzed protein in cells purified from pancreatic cancers (Lau et al., 2020). However, *in vivo* stable isotope tracing has not been performed in somatic stem or progenitor cells, only a few thousand of which can typically be isolated from a mouse. As a result, it is not known how glucose-derived carbon contributes to metabolites in tissue stem cells and restricted progenitor cells *in vivo*. To understand how the utilization of glucose and other nutrients changes during HSC differentiation, we developed an *in vivo* stable isotope tracing analysis method targeting a few metabolites in rare cells. After U^13^C-glucose infusion, bones and thymus were immediately placed on ice, the bones were crushed, cells mechanically dissociated, stained and flow sorted while kept on ice or at 4^0^C as we previously described (Agathocleous et al., 2017) **(Figure 1e).** Metabolites were extracted in 80% acetonitrile and analyzed using HILIC liquid chromatography (DeVilbiss et al., 2021) coupled to a sensitive MS/MS mass spectrometry method. We previously found that in hematopoietic cells, the levels of most metabolites remain stable during bone marrow cell purification performed at cold temperatures (Agathocleous et al., 2017; DeVilbiss et al., 2021). To test if fractional enrichment for metabolites remained stable during hematopoietic cell purification, we compared labeling in metabolites from purified versus freshly isolated bone marrow or thymus cells. Fractional enrichment of U^13^C-glucose derived carbons remained stable during purification of bone marrow and thymus cells in most of the metabolite isotopomers we measured **(Figure 1f-g)**. Changes we observed in some isotopomers after purification may be a consequence of label dilution during the purification process, or the altered cell type composition after purification, for example the loss of stromal cells. 8/10 isotopomers whose enrichment significantly differed between thymus and bone marrow cells before purification, remained significantly different between thymus and bone marrow cells after purification **(Figure 1h)**. Purification did not significantly change the magnitude of labeling differences between thymus and bone marrow for most isotopomers **(Figure 1h)**. Therefore, stable isotope tracing of purified hematopoietic cells measured fractional enrichment differences which largely reflected *in vivo* metabolism. We then performed U^13^C-glucose tracing *in vivo* followed by sorting of hematopoietic cell types. HSCs and MPPs were pooled because they had similarly low levels of glucose uptake **(Figure 1a)** and because they are metabolically more similar to each other than to other cell types (Agathocleous et al., 2017). Glucose-derived labeling on the acetyl group of acetylcarnitine, which originates from acetyl-CoA in the mitochondria (Li et al., 2015), was lower in HSCs/MPPs and in CD48^+^Lineage^-^Sca-1^+^c-Kit^+^ hematopoietic progenitors (HPCs) than in Lineage^-^Sca-1^-^c-Kit^+^ myeloid progenitors (MyP), total sorted bone marrow cells (BM) or total sorted thymus cells **(Figure 1i)**. Carnitine was unlabeled, consistent with acetylcarnitine labeling reflecting labeling of the acetyl group. This result is consistent with the idea that HSCs/MPPs derive very little of their acetyl-CoA from glucose. Aspartate and glutamate, which are directly connected to the TCA cycle via oxaloacetate and α-ketoglutarate respectively **(Figure 1c)**, were labeled to a similarly low or lower level, ∼5-10%, in HSCs/MPPs and HPCs as compared to myeloid progenitors and unfractionated bone marrow cells, while labeling in thymocytes was higher, ∼15-25% **(Figure 1j-k)**. Our results are consistent with the idea that glucose is a major fuel for acetyl groups, aspartate and glutamate in T cell progenitors but not in purified HSCs or restricted myeloid progenitors *in vivo*.

### Pyruvate dehydrogenase is required in double positive thymocytes but not in HSCs or other hematopoietic cells

Glycolysis-derived pyruvate is converted to acetyl-CoA by the pyruvate dehydrogenase (PDH) complex and enters the TCA cycle. To test if PDH is required by hematopoietic cells, we deleted the PDH subunit PDH-E1, encoded by the *Pdha1* gene, in adult hematopoiesis using poly(I:C) treatment of *Mx1Cre;Pdha1^fl/Y^* or *Mx1Cre;Pdha1^fl/fl^* mice (*Pdha1^Δ/Y^* or *Pdha1^Δ/Δ^,* henceforth both denoted as *Pdha1^Δ^*). *Pdha1* deletion in *Pdha1^fl^* mice was previously shown to block PDH complex activity (Johnson et al., 2001; Sidhu et al., 2008). PDH-E1 protein was depleted in *Mx1Cre;Pdha1^Δ^* bone marrow **(Figure S1b)** and HSCs were deleted for *Pdha1* for at least 5 months after poly(I:C) treatment **(Figure S1d)**. The testis-specific homologue *Pdha2* was not expressed in wild type or *Pdha1^Δ^* bone marrow cells **(Figure S1c)**. Young adult *Mx1Cre;Pdha1^Δ^* mice appeared healthy and had normal peripheral white and red blood cell and platelet counts **(Figure S1e-h)**. Bone marrow and spleen cellularity **(Figure 2a-b)**, and the frequency of HSCs, MPPs, restricted progenitors and mature hematopoietic cells in the bone marrow **(Figure 2c, Figure S1i-q)** and spleen **(Figure S2a-j)** did not change after *Pdha1* deletion. The frequency of T cells decreased in the bone marrow and spleen by 10-20 weeks post-deletion **(Figure S1r and S2k)**. Starting at 5 weeks post deletion, the thymus of *Pdha1*-deficient mice was hypocellular **(Figure 2d)**. The number of DP thymocytes, the predominant T cell progenitor in the thymus, and their progeny, the single positive CD4^+^ and CD8^+^ cells, was reduced in *Pdha1^Δ^* mice as compared to littermate controls starting at 5 weeks post-deletion **(Figure 2e-g)**. Consistent with these results, loss of the mitochondrial pyruvate carrier (*Mpc1)* which imports pyruvate in the mitochondria also reduces DP cell number (Ramstead et al., 2020). The number of different subtypes of double negative (DN) cells was unchanged **(Figure 2h, Figure S2o-r)**. The number of CD3^-^CD8^+^ immature single positive (ISP) cells was significantly increased at 20 weeks post deletion, but not at the earlier time of 5 weeks post deletion, when DP cells were affected **(Figure S2s)**. Therefore, PDH deficiency specifically decreased the number of DP thymocytes and their progeny but did not affect other hematopoietic cell types we analyzed.

**Figure 2.**
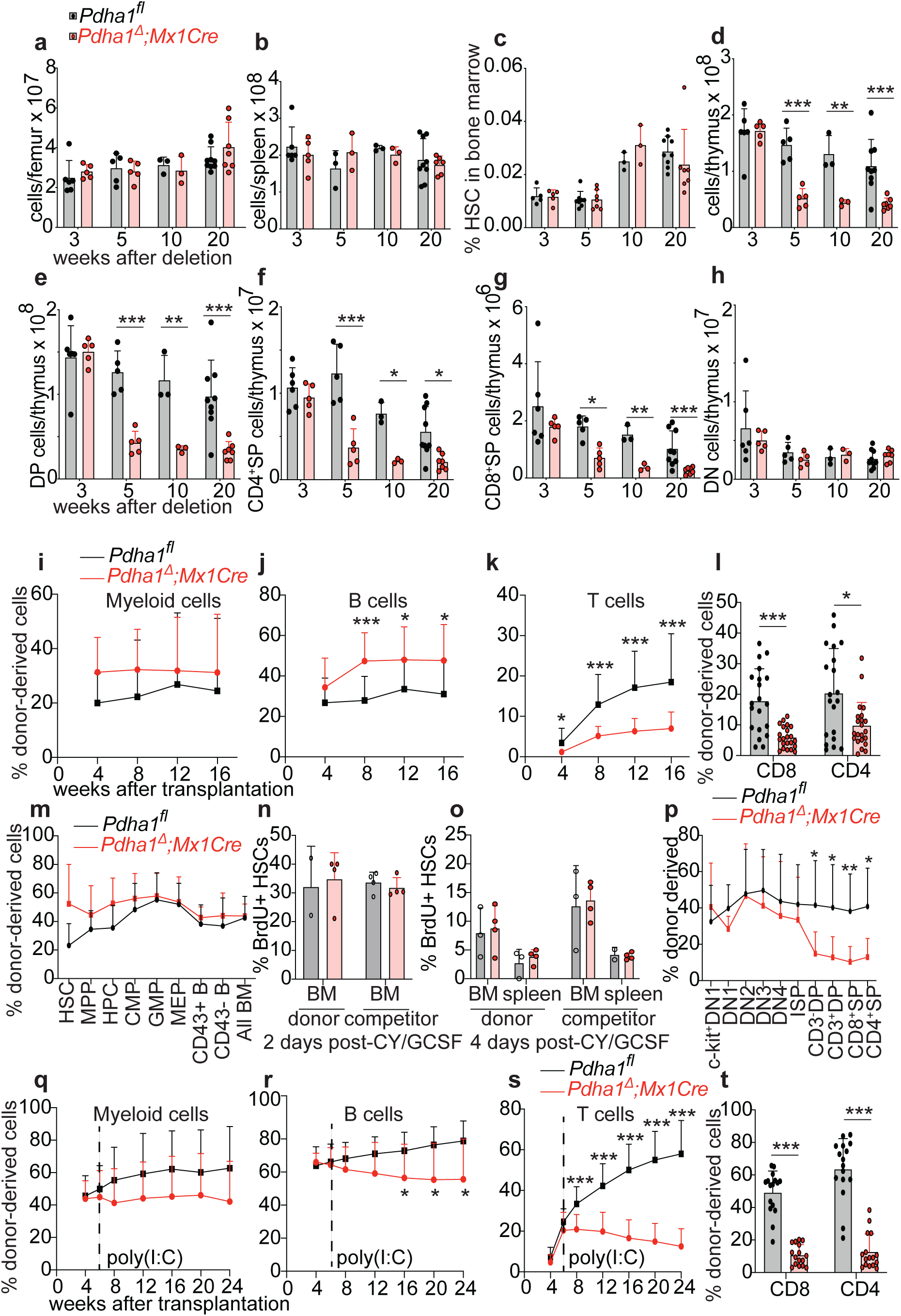
Hematopoietic *Pdha1* deletion impairs the development of DP thymocytes development but not of other hematopoietic cells. **(a-h)** Analysis of hematopoietic cells in the bone marrow, spleen and thymus of *Mx1Cre*;*Pdha1^Δ^* mice and *Pdha1^fl^* littermate controls at the indicated timepoints after poly(I:C)-induced deletion (n=3-9 mice per genotype per timepoint). **(i-p)** Results of competitive transplantation of *Mx1Cre*;*Pdha1^Δ^* or littermate control donor bone marrow cells along with competitor wild-type cells into lethally irradiated wild-type mice. *Pdha1* was deleted with poly(I:C) 3 weeks before transplantation. **(i-l)** Donor cell reconstitution levels in the blood (n=20-24 recipient mice and 5 donor mice/genotype). Myeloid cells were identified as Mac1^+^, B cells as B220^+^, and T cells as CD3^+^. **(l)** Donor reconstitution of CD8^+^SP and CD4^+^SP T cells in the peripheral blood at week 16 (n=20-22 recipient mice). **(m)** The percentage of donor-derived cells in the bone marrow (n=6 mice/genotype). **(n-o)** The proportion of donor and competitor bone marrow cells that incorporated BrdU, 2 and 4 days after cyclophosphamide/G-CSF injection in recipient mice (n=2-4 mice/genotype/timepoint). **(p)** The percentage of donor-derived cells in the thymus (n=10-11 mice/genotype). **(q-t)** Donor cell reconstitution in the blood after competitive transplantation of *Mx1Cre*;*Pdha1^fl^* or littermate control donor bone marrow cells along with competitor wild-type cells into lethally irradiated wild-type mice (a total of 3 donors and 16 recipients per genotype). *Pdha1* was deleted with poly(I:C) 6 weeks after transplantation. **(t)** Donor reconstitution of CD8^+^SP and CD4^+^SP T cells in the peripheral blood was measured at week 24. Timepoints 4 and 6 weeks (before deletion) were excluded from statistical analysis that compared *Mx1Cre*;*Pdha1^fl^* and *Pdha1^fl^* donors. All data represent mean ± s.d. *p < 0.05, **p < 0.01, ***p < 0.001. Statistical significance was assessed with two-way ANOVA (d-g), or a repeated-measures mixed model for data for which some values were missing (i, j, m), or multiple Mann-Whitney tests for data which were not normally distributed (k, l-CD4^+^ cells, t-CD4^+^ cells), t-test with Welch correction (l-CD8^+^ cells, p, t-CD4^+^ cells), t-tests (m) and repeated measures two-way ANOVA with the Geisser-Greenhouse correction (q-s). Multiple comparisons correction was performed by controlling the false discovery rate at 5% or by using multiple comparisons tests as described in the methods.

Proliferating HSCs have been suggested to increase glucose consumption and glucose oxidation compared to quiescent HSCs (Chandel et al., 2016; Liang et al., 2020; Maryanovich et al., 2015; Takubo et al., 2013). To test if glucose oxidation via PDH is required for HSC exit from quiescence and proliferation, we competitively transplanted donor *Pdha1^Δ^* bone marrow cells and wild type competitors into lethally irradiated wild type recipients. *Pdha1* deletion did not significantly change myeloid cell reconstitution, slightly increased B cell reconstitution, and significantly impaired CD4^+^ and CD8^+^ T cell reconstitution **(Figure 2i-l)**. *Pdha1* deletion did not change the proportion of donor-derived HSCs, myeloid or B cell progenitors in the bone marrow of transplant recipient mice **(Figure 2m)**. Administration of cyclophosphamide-GCSF to transplant recipients to induce HSCs to divide (Morrison et al., 1997) showed that *Pdha1* deletion had no significant cell-autonomous effects on the rate of HSC proliferation as measured by 5-bromodeoxyuridine (BrdU) incorporation **(Figure 2n-o)**. These results suggest that oxidation of glucose-derived carbon is not required for HSC maintenance, proliferation, or production of restricted myeloid and B cell progenitors. To understand which cell type is responsible for the T cell development deficit in *Pdha1* deficiency, we analyzed the chimeric thymus of transplant recipients. *Pdha1^Δ^* DN cells or ISPs were found at normal proportions, but *Pdha1^Δ^* DP cells were outcompeted by wild type DP cells **(Figure 2p)**. Therefore, PDH is cell-autonomously required in DP cells. The trend to ISP expansion observed in *Mx1Cre;Pdha1^Δ^* mice **(Figure S2s)** was not observed in chimeric transplant recipients **(Figure 2p)**. This is consistent with a non-cell autonomous effect of PDH deletion on ISPs, possibly secondary to DP depletion, which is known to drive thymic regeneration (Dudakov et al., 2012). The proportion of *Pdha1^Δ^* CD3^+^DP, CD4^+^ or CD8^+^ single positive cells in chimeric recipients was not further reduced than the proportion of *Pdha1^Δ^* CD3^-^DP cells **(Figure 2p, Figure S2t)**, suggesting PDH is mainly required at the CD3^-^DP stage but not at the later thymocyte developmental stages **(Figure S2l)**. Competitive transplantation of *Mx1Cre;Pdha1^fl^* bone marrow into lethally irradiated recipients treated with poly(I:C) 6 weeks after transplantation resulted in a pronounced T cell reconstitution deficit, with no significant changes in myeloid reconstitution and a minor B cell reconstitution deficit **(Figure 2q-t)**. In combination with the results of transplantation after PDH deletion **(Figure 2i-m, p)**, these results suggest PDH is selectively and cell-autonomously required in T cell development but not in HSC, myeloid or B cell development. To test if PDH is required in fetal HSCs, which are highly proliferative, we generated *Vav1Cre;Pdha1^fl^* mice, in which *Pdha1* was deleted during fetal hematopoiesis. Fetal *Pdha1* deletion did not affect the frequency of HSCs, multipotent progenitors, CD48^+^ hematopoietic progenitors or other hematopoietic progenitors or mature cells in the liver at birth, or in the bone marrow and spleen at later timepoints **(Figure S3)**. *Vav1Cre;Pdha1^fl^* mice had reduced thymus cellularity **(Figure S3i, m, s)**, and a decreased number of DP cells and their progeny **(Figure S3l,r,t)**. Therefore, PDH-mediated glucose oxidation was not required by quiescent adult HSCs, adult HSCs induced to proliferate, or proliferating fetal HSCs. Among all hematopoietic cell types we examined, PDH was only required by DP T cell progenitors.

To pinpoint the stage of T cell development that PDH is required for, we generated *CD4Cre;Pdha1^fl^* mice. CD4Cre deletes in DP cells and their progeny T cells (Lee et al., 2001). *CD4Cre;Pdha1^fl^* mice initiated *Pdha1* genomic deletion in CD3^-^DP cells and maintained deletion in their progeny CD3^+^DP, CD3^+^CD4^+^ and CD3^+^CD8^+^ cells but not in their precursor DN or ISP cells as expected **(Figure S4a)**. *CD4Cre;Pdha1^fl^* mice showed no significant changes in thymus cellularity, or the frequency and number of any thymus cell type **(Figure S4b-e)**. To resolve the apparent discrepancy between the cell-autonomous DP loss in *Mx1Cre;Pdha1^fl^* and *Vav1Cre;Pdha1^fl^* mice but not in *CD4Cre;Pdha1^fl^* mice, we blotted for PDH-E1 protein in purified cell types from the *CD4Cre;Pdha1^fl^* thymus. Despite *Pdha1* genomic deletion, CD3^-^DP cells retained some PDH-E1 protein. By the time of DP maturation to the CD3^+^DP stage, PDH-E1 protein was depleted **(Figure S4f)**. Therefore, PDH is likely not required in CD3^+^DP or single positive CD3^+^CD4^+^ and CD3^+^CD8^+^ cells, consistent with our results in transplant recipients that PDH is required in CD3^-^DP cells **(Figure 2p, Figure S2t)**. To confirm that *Pdha1* deletion blocked oxidation of glucose-derived carbon by PDH in *Mx1Cre;Pdha1^fl^* bone marrow and thymocytes but not in *CD4Cre;Pdha1^fl^* thymocytes, we incubated freshly isolated wild type or *Pdha1-*deficient bone marrow cells or purified DP thymocytes in culture with U^13^C-glucose for 2 hours **(Figure S4g)**. *Mx1Cre*-mediated *Pdha1* deletion blocked the flow of labeled carbons from glucose to acetyl-CoA and to citrate in both bone marrow cells and DP thymocytes (more than 90% of which are CD3^-^), suggesting that PDH-E1 protein loss severely reduced or abolished PDH complex activity **(Figure S4h-i)**. In contrast, *CD4Cre*-mediated *Pdha1* deletion did not affect the flow of glucose-derived carbons to acetyl-CoA and citrate in DP thymocytes, consistent with the presence of PDH-E1 protein in *CD4Cre;Pdha1^fl^* CD3^-^DP thymocytes **(Figure S4j-k)**. Therefore, PDH is required for CD3^-^DP cell development or survival but not for the development or survival of downstream or upstream T cell progenitors.

### Initiation of *Pten-*deficient T cell leukemia requires PDH but initiation of *Pten-*deficient myeloid neoplasm does not

To test if PDH is required in leukemia initiation, we crossed *Pdha1^fl^* to *Mx1Cre;Pten^fl^* mice. *PTEN* inactivating mutations are found in 15% of human T cell acute lymphoblastic leukemias (T-ALL) (Liu et al., 2017; Tesio et al., 2017). PTEN is transcriptionally repressed in T-ALL (Palomero et al., 2008) and in some acute myeloid leukemias (AML) (Yoshimi et al., 2011). Pan-hematopoietic deletion of *Pten* using *Mx1Cre* in mice gives rise to T-ALL (Guo et al., 2008; Yilmaz et al., 2006), likely originating from DP thymocytes (Hagenbeek and Spits, 2008), and to myeloid neoplasms, including myeloproliferative neoplasm (Zhang et al., 2006) and AML (Guo et al., 2008; Yilmaz et al., 2006), likely originating from HSCs or restricted myeloid progenitors (Yilmaz et al., 2006) **(Figure 3a)**. *Pdha1* deletion blocked thymomegaly and the increase in the number of DP cells in the thymus of *Pten^Δ/Δ^* mice **(Figure 3b-d)**, without affecting the number of DN thymocytes **(Figure S5a)**. On the contrary, *Pdha1* deletion did not block myeloid expansion in the bone marrow and spleen, splenomegaly, and bone marrow hypocellularity of *Pten^Δ/Δ^* mice **(Figure 3b, e-h)**. *Pten^Δ/Δ^* and *Pten^Δ^*^/*Δ*^*;Pdha1^Δ^*^/*Δ*^ mice had elevated white blood cell counts and *Pten^Δ^*^/*Δ*^*;Pdha1^Δ^*^/*Δ*^ mice had reduced red blood cell counts **(Figure S5b-c)**. Moribund *Mx1Cre;Pten^Δ^*^/*Δ*^ mice developed T-ALL and/or myeloid expansion consistent with a myeloid neoplasm as previously reported (Guo et al., 2008; Hagenbeek and Spits, 2008; Yilmaz et al., 2006; Zhang et al., 2006). Moribund *Mx1Cre;Pten^Δ^*^/*Δ*^*;Pdha1^Δ^*^/*Δ*^ mice harbored myeloid neoplasm, or T-ALL with *Pdha1* alleles which escaped poly I:C-mediated deletion, but we never observed T-ALL with deleted *Pdha1* alleles. To test if *Pdha1* deletion blocked the development of transplantable T-ALL, we transplanted *Pten^Δ/Δ^* or *Pten^Δ/Δ^;Pdha1^Δ^* thymocytes into sublethally irradiated recipients. *Pten^Δ/Δ^* T-ALL was observed in half of the recipient mice, but T-ALL with deleted *Pdha1^Δ^* alleles was not observed in any recipients **(Figure 3i)**.

**Figure 3.**
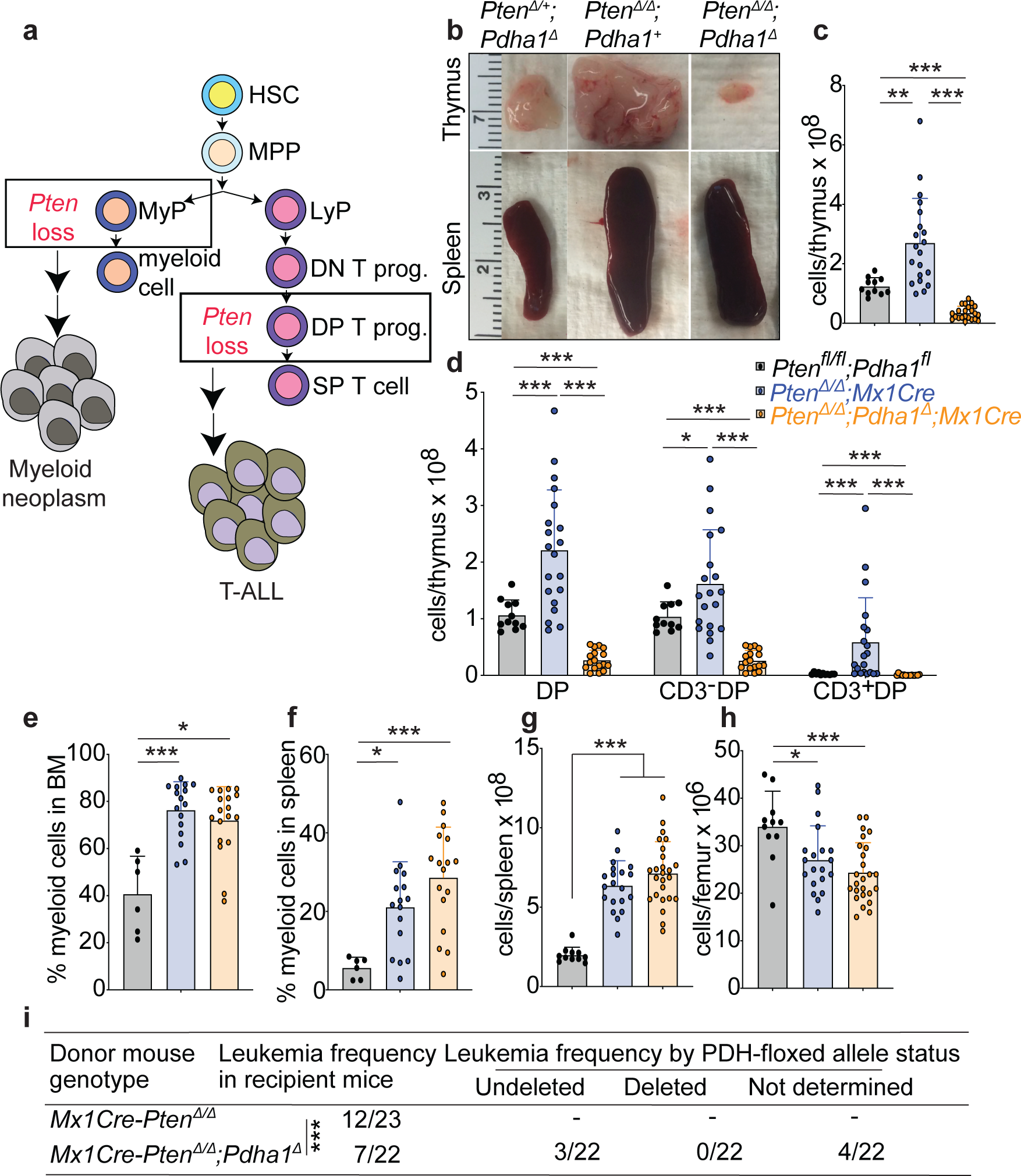
Hematopoietic *Pdha1* deletion inhibits the development of *Pten-*deficient T cell leukemia but not of *Pten-*deficient myeloid neoplasm. **(a)** After *Mx1Cre-*driven hematopoietic *Pten* deletion, *Pten* loss in HSC-derived myeloid progenitors (MyP) gives rise to a myeloid neoplasm, and *Pten* loss in HSC-derived DP T cell progenitors gives rise to T-ALL. Abbreviations: LyP: Lymphoid progenitors; DN, DP T prog.: double negative or double positive T cell progenitors; SP: single positive CD4^+^ or CD8^+^ T cell. **(b-h)** Representative thymus and spleen images, and hematopoietic analysis of mice from the indicated genotypes (n=6-20 mice per genotype). **(i)** Incidence of leukemia in sublethally irradiated mice transplanted with *Mx1Cre;Pten^Δ/Δ^* or *Mx1Cre*;*Pten^Δ/Δ^*;*Pdha1^Δ^* thymocytes. 4 mice which developed leukemia but for which we were unable to establish the *Pdha1* deletion status were excluded from the statistical analysis. Even if all 4 mice turned out to have leukemia with *Pdha1-*deleted alleles, the difference in leukemia incidence between *Mx1Cre;Pten^Δ/Δ^* and *Mx1Cre*-*Pten^Δ/Δ^*;*Pdha1^Δ^* recipients would remain statistically significant. Data represent mean ± s.d. *p < 0.05, **p < 0.01, ***p < 0.001. Statistical significance was assessed with one-way ANOVA with Brown-Forsythe correction (b, c, f) or Kruskal-Wallis test (d) or one-way ANOVA (e,g) or Fisher’s exact test (h). For groups which significantly deviated from normality, a one-way ANOVA test was performed on log-transformed values if they were normally distributed. Multiple comparisons correction was performed by using multiple comparisons tests as described in the methods.

In normal thymus, *Pdha1* deletion increased apoptosis of CD3^+^DP cells, measured by annexin V surface staining 4-6 weeks after deletion, with a trend to increased apoptosis of CD3^-^DP cells, and no effects on the precursor ISP cells **(Figure S5d-f)**. To assess the impact of PDH on proliferation, we administered 5-ethynyl-2-deoxyuridine (EdU) *in vivo* to mark S phase cells. *Pdha1* deletion did not decrease the proportion of EdU-positive DP cells, or their precursor ISP cells **(Figure S5g-i)**. Therefore, PDH is likely required for DP cell survival but not for proliferation, or production from upstream progenitors, consistent with our results from competitive transplantation experiments. *Pdha1* deletion increased apoptosis of *Pten-*deficient CD3^+^DP cells, with a trend to increased apoptosis of *Pten-*deficient CD3^-^DP cells **(Figure S5d-f)**. *Pten-*deficient DP cells were EdU-positive at a significantly higher proportion than wild type DP cells, but *Pten;Pdha1-*deficient DP cells were not **(Figure S5g-i)**, consistent with *Pdha1* deletion preventing the increase in proliferation induced by *Pten* deficiency. These data suggest that PDH deficiency suppresses *Pten-*deficient T-ALL development by inhibiting cell proliferation and promoting apoptosis.

Despite the absence of *Pdha1-*deficient T-ALL, *Mx1Cre;Pten^Δ^*^/*Δ*^*;Pdha1^Δ^*^/*Δ*^ mice nevertheless died at the same time as *Mx1Cre;Pten^Δ^*^/*Δ*^ mice **(Figure 4i)**. This raised the possibility that *Pdha1* deletion may simply delay but not block T-ALL development. To test this, we crossed *Pten^fl^* and *Pten^fl^;Pdha1^fl^* mice with mice carrying *CD2Cre*, which deletes in the lymphoid but not in the myeloid lineage. *CD2Cre;Pten^Δ^*^/*Δ*^ mice had thymomegaly, splenomegaly, liver enlargement, elevated white blood cell counts, bone marrow attrition, thrombocytopenia, expansion of CD3^+^DP pre-leukemic/leukemic cells and T-ALL dissemination to the blood, bone marrow, spleen and liver, all of which were blocked by *Pdha1* deletion **(Figure 4a-h)**. *CD2Cre*-mediated *Pdha1* deletion quadrupled the median survival as compared to deletion of *Pten* alone **(Figure 4i)**. Of the 5 moribund *CD2Cre;Pten^Δ/Δ^;Pdha1^Δ^* mice we analyzed at 10-21 months of age, none had T-ALL, suggesting they died from other causes **(Figure S5j-o)**. PDH is therefore absolutely required for the development of *Pten-*deficient T-ALL. Overall, our results suggest that the requirement for PDH in *Pten-*deficient T-ALL but not *Pten-*deficient myeloid neoplasm mirrors the requirement for PDH in the normal stem or progenitor cell of origin.

**Figure 4.**
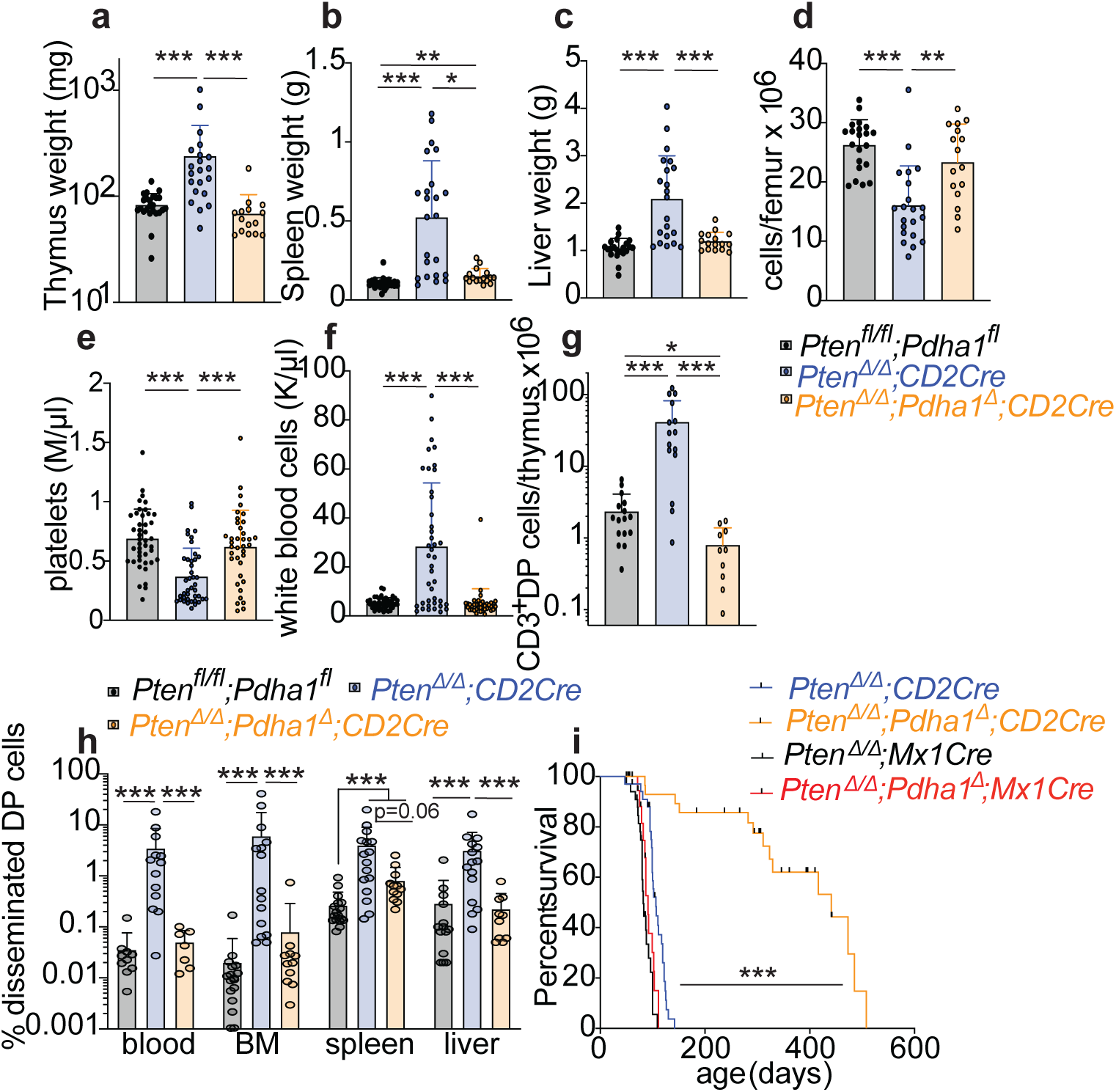
*Pdha1* deletion abolishes the development of *Pten-*deficient T cell leukemia. **(a-d)** Analysis of thymus, spleen, liver and bone marrow of *CD2Cre*;*Pten ^Δ/Δ^*;*Pdha1^Δ^* mice and littermate controls (n=15-21 mice/genotype). **(e-f)** Analysis of peripheral blood in *CD2Cre*;*Pten^Δ/Δ^*;*Pdha1^Δ^* mice and littermate controls (n=37-43 mice/genotype). **(g-h)** Analysis of thymic or disseminated leukemia DP cells in organs (n=7-16 mice/genotype). **(i)** Kaplan-Meier survival curve of mice with hematopoietic (*Mx1Cre-*mediated) or lymphoid (*CD2Cre*-mediated) deletion in *Pten* and *Pdha1* (n=34 mice for *CD2Cre*;*Pten^Δ/Δ^*, n=29 for *CD2Cre*;*Pten ^Δ/Δ^*;*Pdha1^Δ^*, n=33 for *Mx1Cre*;*Pten^Δ/Δ^* and n=23 for *Mx1Cre*; *Pten ^Δ/Δ^*;*Pdha1^Δ^*). All data represent mean ± s.d. *p < 0.05, **p < 0.01, ***p < 0.001. Statistical significance was assessed with Kruskal-Wallis test (a, b, e) or one-way ANOVA with Brown-Forsythe correction (c, h-bone marrow and spleen), or one-way ANOVA (f, g, h-blood and liver), or Mantel-Cox log-rank test (i). For groups which significantly deviated from normality, a t-test or ANOVA test was performed on log-transformed values if they were normally distributed. Multiple comparisons correction was performed by using multiple comparisons tests as described in the methods.

### Glycolysis and the TCA cycle are connected through PDH in the thymus but not in the bone marrow

To understand the metabolic consequences of *Pdha1* deletion, we infused wild type and *Mx1Cre;Pdha1^Δ^* mice with U^13^C-glucose and traced its contribution to glycolytic and TCA cycle metabolites. The first turn of the TCA cycle after entry of U^13^C-glucose-derived acetyl-CoA through the PDH reaction yields metabolites labeled with 2 heavy carbons (M+2) **(Figure 5a)**. PDH deletion in the bone marrow did not significantly reduce the fractions of M+2 labeled TCA cycle metabolites citrate, succinate, fumarate, and malate. PDH deletion in the bone marrow also did not significantly reduce the fractions of M+2 labeled aspartate, glutamate and glutamine, metabolites that can be derived from, or feed into the TCA cycle **(Figure 5b)**. PDH deletion did not affect the pyruvate(M+3)/citrate(M+2) ratio in the bone marrow *in vivo* **(Figure 5d)**. These results suggest that even the low levels of glucose-derived carbon labeling observed in the TCA cycle in the bone marrow **(Figure 1d)** are not derived from cell-autonomous PDH reaction but indirectly from other circulating nutrients which are labeled from infused U^13^C-glucose in other tissues. By contrast, PDH deletion significantly reduced the levels of M+2 labeled citrate, succinate, malate, aspartate and glutamate in the thymus of the same animals **(Figure 5c)**. Because DP cells comprise the great majority of thymocytes, these results likely reflect glucose catabolism in DP cells. Therefore, a significant fraction of the glucose-derived carbons in the TCA cycle are derived from the PDH reaction in thymus but not in bone marrow. These *in vivo* tracing results contrast with the results of *ex vivo* tracing of freshly isolated cells, which showed that both bone marrow and DP thymocytes perform PDH-mediated glucose oxidation **(Figure S4g-i)**. Therefore, standard *ex vivo* culture is likely unsuitable to delineate *in vivo* hematopoietic cell metabolism, stressing the need for developing *in vivo* approaches.

**Figure 5.**
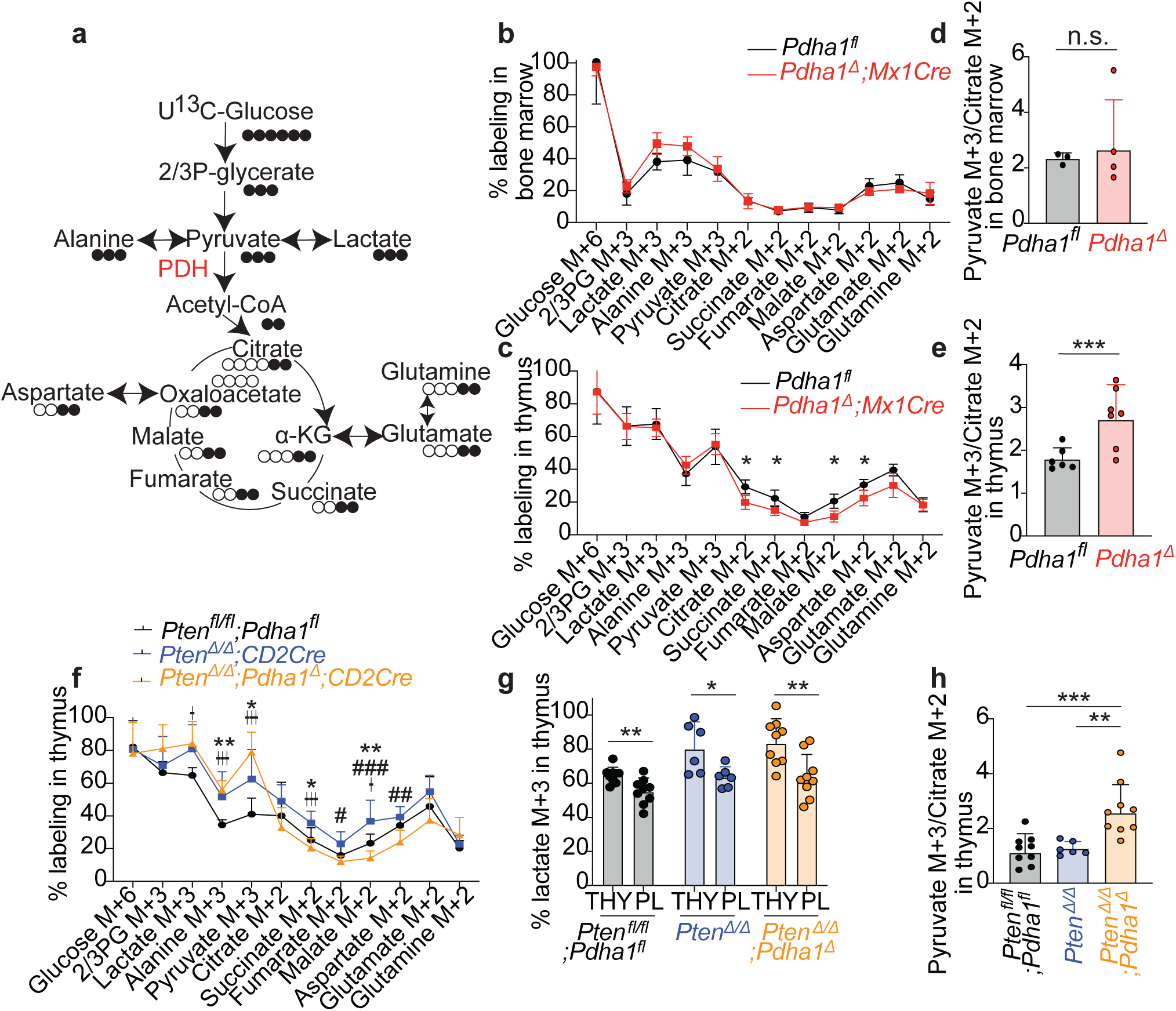
Glycolysis is connected to the TCA cycle through PDH in the thymus but not in the bone marrow. **(a-e)** *In vivo* U^13^C-glucose infusion in *Mx1Cre*;*Pdha1^Δ^* and littermate controls (n=3-7 mice/genotype. Data shown in Fig. 1d, which compares glucose utilization in wild type thymus and bone marrow, are derived from this experiment). **(a)** Schematic depicting expected metabolite labeling patterns in glycolysis and after the first turn of the TCA cycle. **(b, c)** Metabolite labeling in the bone marrow and thymus. **(d-e)** Pyruvate (M+3)/Citrate (M+2) ratio in the bone marrow and thymus. **(f-h)** *In vivo* U^13^C-glucose infusion in *CD2Cre*;*Pten^Δ/Δ^;Pdha1^Δ^* mice and littermate controls (n=6-9 mice/genotype). **(f)** Metabolite labeling in the thymus. * compares wild-type *vs CD2Cre*;*Pten^Δ/Δ^,* † compares wild-type *vs CD2Cre*;*Pten^Δ/Δ^;Pdha1^Δ^,* and # compares *CD2Cre*;*Pten^Δ/Δ^ vs CD2Cre*;*Pten^Δ/Δ^;Pdha1^Δ^* mice. **(g)** Comparison of % lactate M+3 labeling in thymus and blood. **(h)** Pyruvate (M+3)/Citrate (M+2) ratio in the thymus. All data represent mean ± s.d. or for (d,e,h) geometric mean ± geometric s.d. *p < 0.05, **p < 0.01, ***p < 0.001. Statistical significance was assessed with t-tests (b, except for glutamate, c except for glutamine, d, e), one-way ANOVA (f for 3-phosphoglycerate, alanine, fumarate, glucose, malate, pyruvate, succinate, glutamate, citrate), Kruskal-Wallis test (f for aspartate, glutamine), one-way ANOVA with Brown-Forsythe correction (f for lactate, h), matched t-tests (g) or Mann-Whitney tests (b for glutamate, c for glutamine). For groups which significantly deviated from normality, a t-test or ANOVA test was performed on log-transformed values if they were normally distributed. Multiple comparisons correction was performed by controlling the false discovery rate at 5% or by using multiple comparisons tests as described in the methods.

To test how leukemic transformation impacts glucose catabolism, we measured the contribution of U^13^C-glucose to metabolites in wild type, *Pten-*deficient, and *Pten;Pdha1* deficient thymus. We took advantage of the fact that DP cells are the predominant cell type in the thymus in all our genetic backgrounds, therefore metabolic changes observed in the thymus likely reflect metabolic changes in DP cells. We used our *CD2Cre*-mediated deletion model because cells in *Mx1Cre;Pten^fl/fl^;Pdha1^fl^* mice often escape poly I:C-mediated *Pdha1* deletion due to selective pressure against PDH loss*. Pten* deletion significantly increased the contribution of glucose-derived carbon to pyruvate, lactate, alanine, succinate and malate **(Figure 5f)**, consistent with the well-known effects of PI3K pathway activation in stimulating glucose use in multiple cell types (Elstrom et al., 2004), including in T-ALL cells *in vitro* (Herranz et al., 2015). The fraction of glucose-derived lactate was higher in the thymus than in the plasma of all genotypes **(Figure 5g)**, consistent with intracellular lactate production from glucose rather than lactate import from the blood, unlike in other tissues or cancers (Faubert et al., 2017; Hui et al., 2017). These results suggest that the main effect of *Pten* deletion on thymus central carbon metabolism is to increase glycolytic production of pyruvate, without an additional increase in glucose-derived contribution to the TCA cycle. *Pdha1* deletion in *Pten-*deficient thymus maintained the high labeling of glycolytic metabolites from glucose and further increased the contribution of glucose to pyruvate. *Pdha1* deletion significantly reduced the contribution of glucose-derived carbon to TCA cycle metabolites but did not abolish it. Therefore, some glucose-derived carbon input to the TCA cycle is derived from pyruvate through the PDH reaction, and some is derived from other circulating metabolites labeled from glucose, such as glutamine (Herranz et al., 2015) or acetate (Jakkamsetti et al., 2019). *Pdha1* deletion increased the pyruvate(M+3)/citrate(M+2) ratio in the thymus, consistent with a block of the PDH reaction **(Figure 5e, h)**. These results raise the question of why PDH, which facilitates entry of glucose-derived carbon into the TCA cycle, is nevertheless required in *Pten-*deficient thymocytes, which have higher glycolysis than normal thymus or bone marrow.

### PDH regulates pyruvate levels, glutathione, and redox homeostasis in the thymus

To understand the metabolic consequences of PDH deficiency, we performed metabolomics in wild type and *Pdha1-*deficient bone marrow and thymus. *Pdha1* deficiency did not significantly change the levels of any metabolite in the bone marrow, out of a total of 223 detected metabolites, providing further evidence for the functional disconnection of glycolysis and the TCA cycle in the marrow **(Figure 6a)**. In contrast, *Pdha1* deficiency significantly changed the levels of 42 out of 242 detected metabolites in the thymus **(Figure 6b, Supplementary Table 1)**. *Pdha1*-deficient thymus had 10-fold increased level of pyruvate in the thymus but not in the bone marrow **(Fig 6c, and Figure S6a)**. Surprisingly, *Pdha1* deficiency did not change the levels of acetyl-CoA or the levels of multiple TCA cycle intermediates including citrate/isocitrate, α-ketoglutarate, malate, succinate and fumarate in the thymus **(Figure 6d)**. The acetylcarnitine:carnitine ratio was significantly reduced in *Pdha1-*deficient thymus, consistent with channeling of PDH-derived acetyl-CoA to carnitine acetyltransferase (Li et al., 2015), but acetylation of several other metabolites did not significantly change **(Figure S6b)**. The levels of many long and medium chain carnitines were increased in *Pdha1* deficient thymus, suggesting compensatory changes in oxidation of other substrates replaced PDH-derived acetyl-CoA **(Figure S6c)**. Mitochondrial mass and membrane potential were unaltered in *Pdha1-*deficient thymocytes and in other hematopoietic cells **(Figure S6d-l)**. These results suggest that even though PDH in the thymus delivers glucose-derived carbons to the TCA cycle, in its absence other fuels can generate sufficient acetyl-CoA to sustain TCA cycle metabolite levels, mitochondrial activity, and acetylation reactions. It remains possible that acetyl-CoA dependent metabolic pathways other than the ones we examined are sensitive to loss of PDH-mediated acetyl-CoA production.

**Figure 6.**
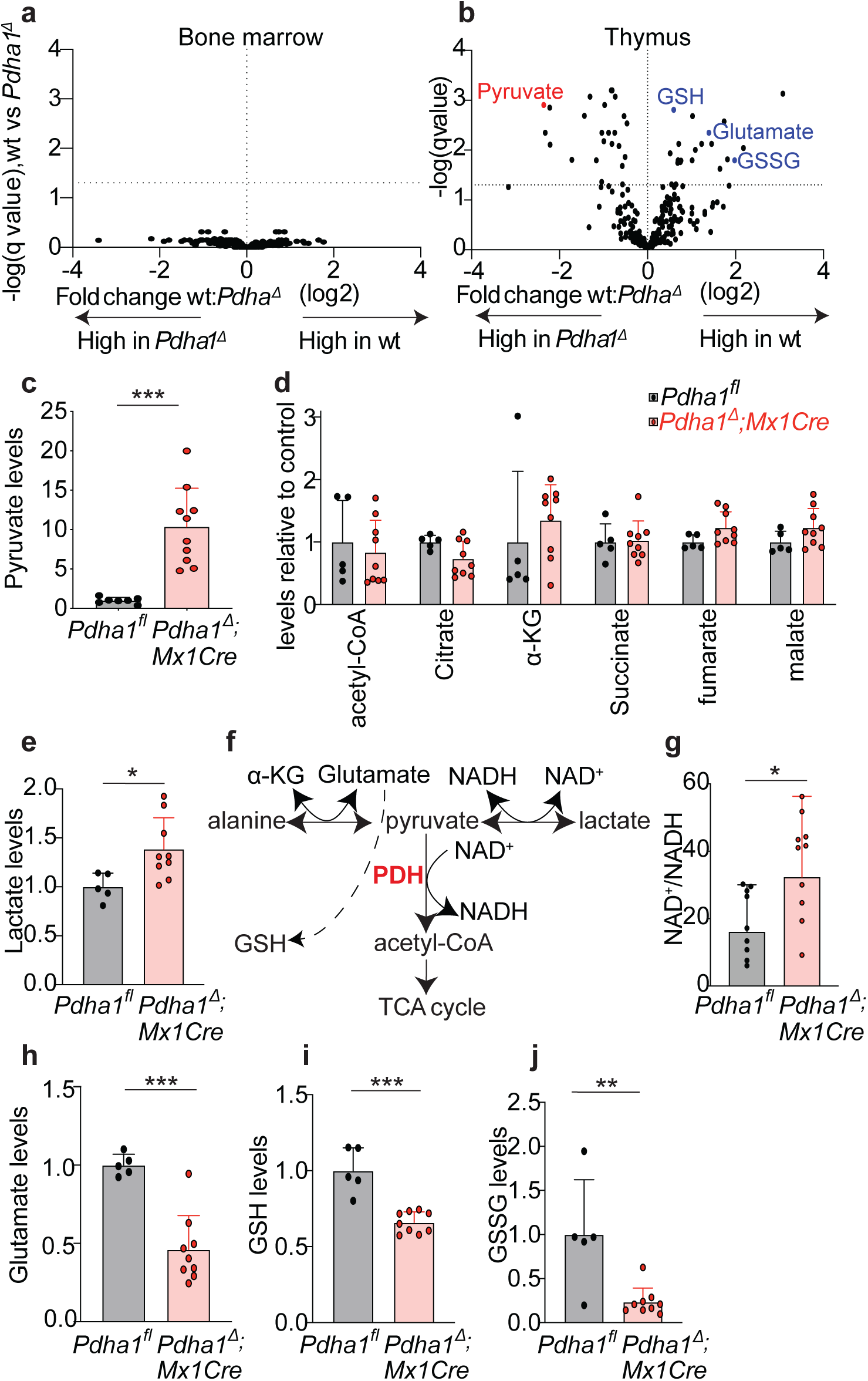
*Pdha1* deletion in the thymus causes accumulation of pyruvate and aberrant redox balance. **(a-b)** Volcano plots showing results of metabolomics analysis in the bone marrow and thymus of *Mx1Cre*;*Pdha1^Δ^* mice and *Pdha1^fl^* (wild type) littermate controls. Each dot represents a metabolite (analysis from n=5 *Pdha1^fl^* and 9 *Pdha1^Δ^* mice). The q value represents the p value of comparing the mean of each metabolite in the two treatments, after multiple comparisons correction. **(c)** Pyruvate levels in the thymus measured with o-phenylenediamine derivatization and normalized to the average of *Pdha1^fl^* samples (n=7-10 mice/genotype). **(d, e)** Acetyl-CoA, TCA cycle metabolites and lactate levels in the thymus, normalized to the average of *Pdha1^fl^* samples. **(f)** The connection of the PDH reaction to pyruvate, lactate, NAD^+^/NADH, glutamate and glutathione. **(g)** The ratio of NAD^+^/NADH in the thymus of *Mx1Cre*;*Pdha1^Δ^* mice and *Pdha1^fl^* mice (n=9-10 mice/genotype). **(h-j)** Levels of glutamate and reduced (GSH) and oxidized (GSSG) glutathione in the thymus (n=5-9 mice/genotype). All data represent mean ± s.d. or for (g) geometric mean ± geometric s.d. *p < 0.05, **p < 0.01, ***p < 0.001. Statistical significance was assessed with t-tests (a, b, c, e, g) or Mann-Whitney tests (c for fumarate and α-KG). For data which significantly deviated from normality, a t-test was performed on log-transformed values if they were normally distributed. Multiple comparisons correction was performed by controlling the false discovery rate at 5% or by using multiple comparisons tests as described in the methods.

Consistent with a block in pyruvate utilization, lactate levels were also increased in *Pdha1*-deficient thymus, albeit to a lesser extent than the increase in pyruvate levels **(Figure 6e)**. The pyruvate:lactate ratio is proportional to the cytosolic NAD^+^:NADH ratio (Williamson et al., 1967) **(Figure 6f)**. In *Pdha1-*deficient thymus, the NAD^+^:NADH ratio was elevated by 2-fold compared to the wild type thymus, consistent with the increase in the pyruvate:lactate ratio **(Figure 6g)**. This is consistent with a recent report that PDH activation decreases the NAD^+^:NADH ratio (Luengo et al., 2020). *Pdha1*-deficient thymus had decreased levels of glutamate **(Figure 6h)**. A reduction in glutamate levels might reflect reduced production from the TCA cycle, however α-ketoglutarate, the TCA cycle glutamate precursor, was not depleted in *Pdha1*-deficient thymus **(Figure 6d)**. These results are more consistent with elevated pyruvate or NAD^+^/NADH in *Pdha1-*deficient thymus causing glutamate depletion through transamination reactions, many of which use glutamate, rather than with decreased TCA cycle-mediated glutamate production. Reduced and oxidized glutathione (GSH and GSSG), which rely on glutamate for biosynthesis, were also depleted in *Pdha1-*deficient thymus **(Figure 6i-j)**. This suggests that a major role of PDH in the thymus is to metabolize pyruvate and maintain redox homeostasis, glutamate levels, and glutathione synthesis.

To understand how the metabolic consequences of *Pdha1* deficiency in the normal thymus compare to those in the setting of *Pten-*deficiency, we compared *Pten^fl/fl^;Pdha1^fl^* (wild type), *CD2Cre;Pten^Δ/Δ^* and *CD2Cre;Pten^Δ/Δ^;Pdha1^Δ^* thymus. The thymus from each genotype was metabolically distinct suggesting significant metabolic reprogramming after *Pten* or *Pten;Pdha1* deletion **(Figure 7a, Figure S7a)**. *Pten^Δ/Δ^;Pdha1^Δ^* thymus was more metabolically similar to wild type than to *Pten^Δ/Δ^* thymus, suggesting *Pdha1* deletion reversed many metabolic changes caused by *Pten* deletion **(Figure 7a-b)**. PDH deletion in *Pten^Δ/Δ^* thymus had metabolic consequences that partly overlapped with those caused by PDH deletion in normal thymus **(Figure S7b, Supplementary Table 2)**. Similar to the wild type thymus, PDH deletion in the *Pten^Δ/Δ^* thymus increased the levels of pyruvate and lactate, increased the NAD^+^/NADH ratio, decreased glutamate and glutathione levels, and did not affect levels of acetyl-CoA or the TCA cycle metabolites citrate, α-ketoglutarate, succinate and malate **(Figure 7c-i)**. The total levels of histone H3K27, H3K9 and H4K16 acetylation did not significantly change in *Pten^Δ/Δ^*;*Pdha1^Δ^* as compared to *Pten^Δ/Δ^* or wild type thymus **(Figure S7c-d)**. This data suggests that histone acetylation in *Pten^Δ/Δ^* thymus does not depend on PDH activity though it does not exclude the possibility that histone acetyl marks other than the ones we examined, or marks on specific loci, are regulated by PDH. Mitochondrial mass was unchanged in *Pten^Δ/Δ^;Pdha1^Δ/Δ^* as compared to *Pten^Δ/Δ^* or wild type thymus **(Figure S7e)**. Our results suggest that PDH is not required in *Pten-* deficient thymus to maintain acetyl-CoA levels, TCA cycle metabolites or histone acetylation, but is required to dispose of glucose-derived pyruvate, maintain redox homeostasis and support glutamate levels and glutathione synthesis.

**Figure 7.**
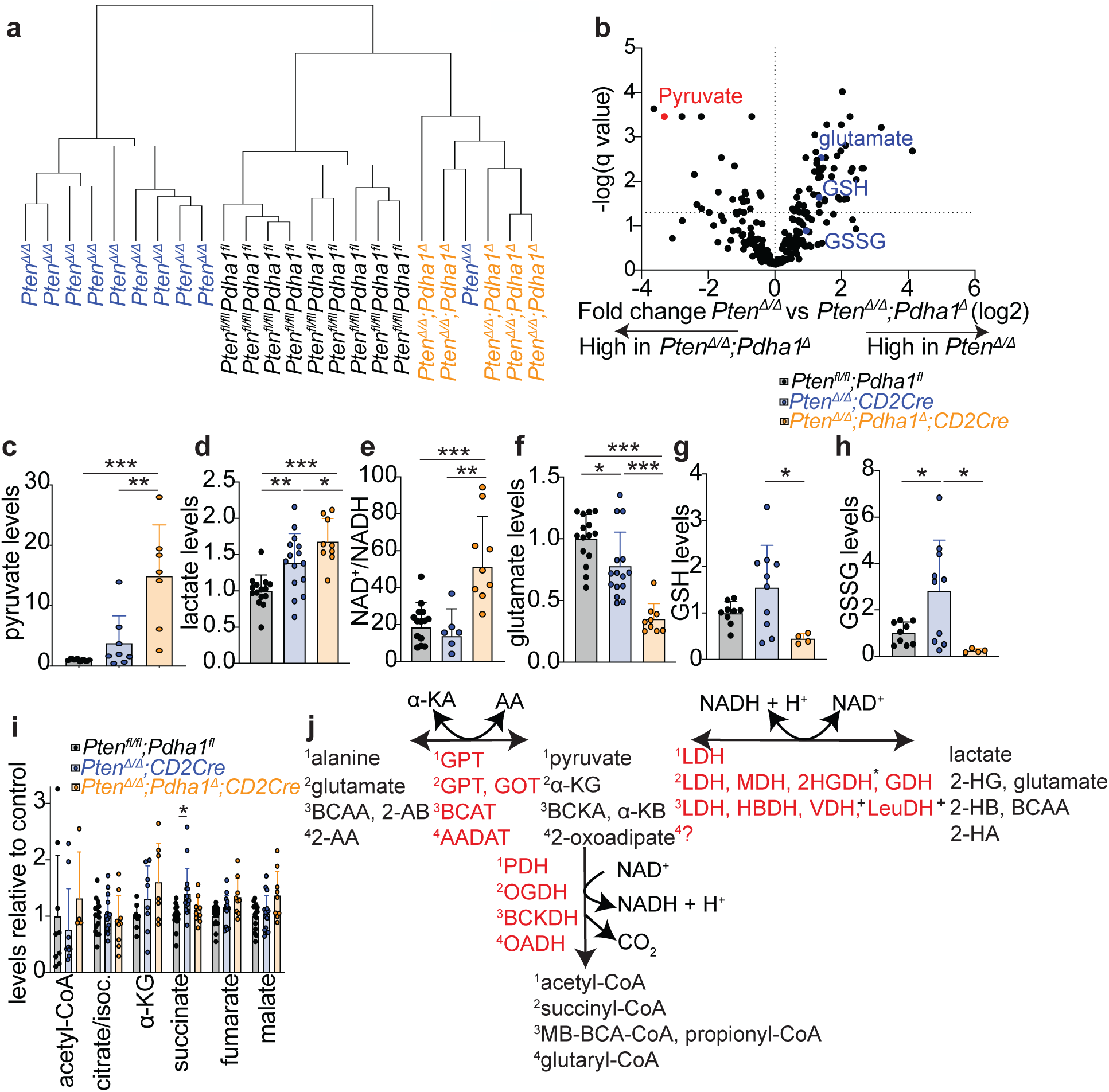
*Pdha1* deletion prevents *Pten-*deficiency induced metabolic reprogramming in the thymus, and causes accumulation of pyruvate and aberrant redox balance. **(a)** Unsupervised clustering of metabolomic analysis from the thymus of *CD2Cre*;*Pten^Δ/Δ^;Pdha1^Δ^* mice and littermate controls (n=5-10 mice/genotype). Clustering was performed using the Euclidean distance measure with Ward clustering algorithm. **(b)** Volcano plot comparing the metabolome of *Pten^Δ/Δ^* and *Pten^Δ/Δ^*;*Pdha1^Δ/Δ^* mice. Each dot represents a metabolite. The q value represents the p value of comparing the mean of each metabolite in the two treatments, after multiple comparisons correction (n=5-10 mice/genotype). **(c)** Pyruvate levels in the thymus measured with o-phenylenediamine derivatization and normalized to the average of *Pten^fl/fl^;Pdha1^fl^* samples (n=7-8 mice/genotype). **(d, f)** Levels of lactate and glutamate in the thymus (n=9-15 mice/genotype). **(e)** The ratio of NAD^+^/NADH in the thymus (n=6-14 mice/genotype). **(g-h)** Levels of reduced (GSH) and oxidized (GSSG) glutathione in the thymus measured using formic acid extraction as described in the methods (n=4-10 mice/genotype). **(i)** Levels of TCA cycle metabolites in the thymus. **(j)** Schematic of coupling between α-ketoacids and redox. α-ketoacids can either act as an electron donor through their respective α-ketoacid dehydrogenases (PDH, OGDH, BCKDH, OADH) or as an electron acceptor through an NADH-dependent dehydrogenase, or as a substrate for a transamination reaction. Loss of PDH in the thymus converts pyruvate from an electron donor to an electron acceptor in the LDH reaction, and promotes its transamination with glutamate depletion in the GPT reaction. The schematic represents parallel reactions, numbered 1-4, for different α-ketoacid dehydrogenases. Enzymes are shown in red. Abbreviations: LDΗ lactate dehydrogenase, MDH malate dehydrogenase, 2HGDH 2-hydroxyglutarate dehydrogenase, GDH glutamate dehydrogenase, HBDH α-hydroxybutyrate dehydrogenase, VDH Valine dehydrogenase, LeuDH Leucine dehydrogenase, GPT alanine aminotransferase, GOT aspartate aminotransferase, BCAT branched chain amino acid aminotransferase, AADAT aminoadipate aminotransferase. α-ΚΑ α-ketoacid, AA amino acid, BCKA branched-chain keto acid, α-KB α-ketobutyrate, 2-AB 2-aminobutanoate, 2-AA 2-aminoadipate, 2-HG 2-hydroxyglutarate, 2-HB 2-hydroxybyturate, BCAA branched chain amino acid, 2-HA 2-hydroxyadipate, BCA-CoA branched chain acyl-CoA. + LeuDH and VDH are bacterial enzymes, *2HGDH is FAD-dependent. Some reactions involve additional substrates/products, such as ammonia. All data represent mean ± s.d. or for (e) geometric mean ± geometric s.d. *p < 0.05, **p < 0.01, ***p < 0.001. Statistical significance was assessed with multiple t-tests (b) or one-way ANOVA (c-h). For data which significantly deviated from normality, a one-way ANOVA was performed on log-transformed values if they were normally distributed. Multiple comparisons correction was performed by controlling the false discovery rate at 5% or by using multiple comparisons tests as described in the methods.

## DISCUSSION

90% of cells in the human body are produced by the bone marrow, and most of them live only a few days to a few months before being replaced, illustrating the marrow’s enormous proliferative activity at homeostasis, which exceeds that of all other human tissues (Sender et al., 2016; Sender and Milo, 2021). Yet we find that in this tissue, glycolysis and the TCA cycle were disconnected. Little or none of glucose-derived carbon entered the TCA cycle through the PDH reaction, and PDH deletion did not affect the function of any of the marrow populations we examined, including HSCs and restricted myeloid, B and erythroid progenitors. This suggests that nutrients other than glucose, including glutamine or fatty acids (Ito et al., 2012; Oburoglu et al., 2014; Rajagopalan et al., 2015) may generate acetyl-CoA and fuel the TCA cycle in HSCs and restricted hematopoietic progenitors.

The metabolic consequences of PDH deficiency in the thymus and T-ALL were surprising. The main function of PDH, including in cancer cells, is thought to be the synthesis of acetyl-CoA for the TCA cycle, lipid synthesis and other acetylation reactions, including acetylation of histones (Chen et al., 2018; Nagaraj et al., 2017; Sutendra et al., 2014). We find that PDH was instead required in the thymus to regulate the NAD^+^/NADH ratio, pyruvate and glutamate levels. PDH deficiency in the brain also depletes glutamate levels without depleting TCA cycle metabolites (Jakkamsetti et al., 2019), suggesting that the metabolic role of PDH we described in the thymus may be present in other glucose-avid tissues. In the absence of PDH, pyruvate can be transaminated using glutamate, a redox-neutral reaction which produces α-ketoglutarate and alanine, or can act as an electron acceptor in the lactate dehydrogenase (LDH) reaction, oxidizing NADH to NAD^+^. PDH converts pyruvate from an electron acceptor to an electron donor, reducing NAD^+^ to NADH, accompanied by oxidative decarboxylation to acetyl-CoA. The other known α-keto acid dehydrogenases also convert their α-keto acid substrate from an electron acceptor that produces NAD^+^, to an electron donor that produces NADH, accompanied by oxidative decarboxylation to form an acyl thioester linkage with coenzyme A **(Figure 7j)**. It is possible that in tissues with high rates of uptake or production of the corresponding α-keto acid, α-keto acid dehydrogenases are required for redox, α-keto acid and amino acid homeostasis, rather than for nutrient catabolism.

The inference that cancer cells are metabolically different from normal cells in the same tissue is largely drawn from measurements comparing metabolism in cancer cells to the corresponding bulk normal tissue. Such comparisons cannot distinguish between metabolic differences which are caused by oncogenic mutations and those which are inherited by the specific cell type that the cancer originates from and maintained by oncogenic mutations. We observed multiple changes in metabolite levels in *Pten-*deficient compared to wild type thymus, despite the fact that both are composed predominantly of DP cells, consistent with *Pten-*deletion driven metabolic reprogramming. We found that a major effect of *Pten* deletion was to accentuate the pre-existing propensity of DP cells to use and rely on glucose oxidation. Importantly, our results show that for *Pten-*driven leukemia cells, their dependence on glucose oxidation is a function of their normal progenitor cell of origin and not of the cancer driver mutation. It will be interesting to ask which of the metabolic features of cancers from other tissues arise during transformation, and which reflect the metabolism of their normal stem or progenitor cell of origin.

## Acknowledgements

M.A. is a Cancer Prevention and Research Institute of Texas Scholar and an American Society of Hematology Faculty Scholar. We thank S. Morrison and R. DeBerardinis for discussions and B. Harris for comments on the manuscript. We thank A. Tasdogan, B. Faubert and J. Sudderth for help with glucose infusions and GC-MS analysis. We thank L. Nguyen and J. Rose III for mouse colony management, N. Loof and the Moody Foundation Flow Cytometry Facility for flow cytometry and L. Zacharias and the CRI Metabolomics Facility for assistance with metabolomics. This work was supported by the Cancer Prevention and Research Institute of Texas (RR180007), Alex’s Lemonade Stand Foundation for Childhood Cancer and the American Society of Hematology.

## Authors’ Contributions

S.J., S.M. and M.A. designed and performed all experiments and analyzed all data. S.M., T.P.M., M.S.M.S. and M.A. developed metabolomics methods. Z.Z. helped with statistical analysis. E.P. helped with histone acetylation experiments. M.A. wrote the manuscript with help from S.J.

## Methods

### Mice

*Pten*^fl/fl^ (Groszer et al., 2001), *Pdha1*^fl/fl^ (Johnson et al., 2001), *Mx1-Cre* (Kühn et al., 1995), *Vav1-Cre(Ogilvy et al., 1998), CD2-Cre* (Zhumabekov et al., 1995), and *CD4-Cre* (Lee et al., 2001) mice were previously described. Mice were on a C57BL background. Both male and female mice were used in all studies. *Pdha1* is an X-linked gene, and male mice are hemizygous for *Pdha1*. In the figures and text, male *Pdha1^fl/Y^* and female *Pdha1^fl/fl^* mice are both denoted as *Pdha1^fl^*. Μale *Pdha1^Δ/Y^* and female *Pdha1^Δ/Δ^* mice are both denoted as *Pdha1^Δ^*. We did not identify differences in our assays between *Pten^Δ/Δ^;Pdha1^Δ/+^* and *Pten^Δ/Δ^* mice, therefore in most experiments *Pten^Δ/Δ^;Pdha1^Δ/+^* female mice and *Pten^Δ/Δ^;Pdha1^Y/+^* male mice were included in the same category and denoted as ‘*Pten^Δ/Δ’^*. ‘Wild-type’ littermate controls in experiments involving *Pten* deletion included Cre-negative *Pten^fl/fl^;Pdha1^fl/+^* females, *Pten^fl/fl^;Pdha1^fl/fl^* females, *Pten^fl/fl^;Pdha1^+/Y^* males and *Pten^fl/fl^;Pdha1^fl/Y^* males. In the figures and text, these mice are denoted as *Pten^fl/fl^;Pdha1^fl^.* In *Mx1Cre* models, mice were injected with seven intraperitoneal injections of 60 μg poly(I:C) (GE Healthcare) every other day at 4-6 weeks of age to induce deletion. C57BL/Ka-Thy-1.1 (CD45.2) and C57BL/Ka-Thy-1.2 (CD45.1) mice were used for transplantation experiments. For cyclophosphamide/G-CSF induction of HSC proliferation, mice were injected with 4 mg of cyclophosphamide i.p. followed by 2 or 4 daily s.q. injections of 5 μg granulocyte colony-stimulating factor (G-CSF). Mice were housed in the Animal Resource Center at the University of Texas Southwestern Medical Center and all procedures were approved by the UT Southwestern Institutional Animal Care and Use Committee.

### Cell isolation for experiments other than metabolomics

Cell populations were defined with the following markers: CD150^+^CD48^-^Lineage^-^Sca-1^+^c-Kit^+^ hematopoietic stem cells (HSCs), CD150^-^CD48^-^Lineage^-^Sca-1^+^c-Kit^+^ multipotent progenitor cells (MPPs), CD48^+^Lineage^-^Sca-1^+^c-Kit^+^ hematopoietic progenitor cells (HPCs), CD150^-^CD48^+^Lineage^−^Sca-1^+^c-Kit^+^ (HPC1), CD150^+^CD48^+^Lineage^−^Sca-1^+^c-Kit^+^ (HPC2), CD34^+^CD16/32^-^Lineage^-^Sca-1^-^c-Kit^+^ common myeloid progenitors (CMPs), CD34^+^CD16/32^+^Lineage^-^Sca-1^-^c-Kit^+^ granulocyte–macrophage progenitors (GMPs), CD34^-^CD16/32^-^Lineage^-^Sca-1^-^c-Kit^+^ megakaryocyte–erythroid progenitors (MEPs), Mac-1^+^Gr-1^+^ and Mac-1^+^Gr-1^-^ myeloid cells, CD71^mid^Ter-119^-^ (R0), CD71^+^Ter-119^-/mid^ (R1), CD71^+^Ter-119^+^ (R2), CD71^mid^Ter-119^+^ (R3), CD71^-^Ter-119^+^ (R4) erythroid cells, CD3^+^ T cells, and B220^+^ B cells. The Lineage antibody cocktail for HSCs and bone marrow progenitors consisted of CD2, CD3, CD5, CD8, Ter-119, B220 and Gr-1 antibodies. T cell progenitor populations in the thymus were defined with the following markers, after excluding Mac-1^+^ or Ter-119^+^ or B220^+^ cells: CD3^+^CD4^+^CD8^-^ (CD4^+^ single positive, SP), CD3^+^CD4^-^CD8^+^ (CD8^+^ SP), CD3^-^CD4^-^CD8^+^ (ISPs), CD3^-^CD4^+^CD8^+^ (CD3^-^DPs), CD3^+^CD4^+^CD8^+^ (CD3^+^DPs), CD4^-^CD8^-^ (DNs), CD4^-^CD8^-^CD44^+^CD25^-^ (DN1), CD4^-^CD8^-^CD44^+^CD25^+^ (DN2), CD4^-^CD8^-^CD44^-^ CD25^+^ (DN3), CD4^-^CD8^-^CD44^-^CD25^-^ (DN4). Analysis and sorting were performed using a FACSAria flow cytometer (BD Biosciences) or a BD FACSCanto (BD Biosciences). Data were analysed using FACSDiva (BD Biosciences).

| Name | Clone | Supplier | Catalog # |
| --- | --- | --- | --- |
| CD150 | TC15-12F12.2 | BioLegend | 115904 |
| CD48 | HM48-1 | BioLegend | 103426 |
| Sca1 | D7 | BioLegend | 122514 |
| CD117/cKit | 2B8 | eBioscience | 47-1171-82 |
| CD34 | RAM34 | eBioscience | 50-0341-82 |
| CD16/32 | 93 | BioLegend | 101333 |
| CD11b | M1/70 | eBioscience | 17-0112-83 |
| Gr-1 | RB6-8C5 | BioLegend | 108406 |
| CD71 | R17217 | eBioscience | 12-0711-83 |
| Ter119 | TER-119 | BioLegend | 116206 |
| CD3ε | 17A2 | TONBO biosciences | 35-0032 |
| CD45R/B220 | RA3-6B2 | TONBO biosciences | 35-0452 |
| CD2 | RM2-5 | TONBO biosciences | 35-0021 |
| CD5 | 53- 7.3 | BioLegend | 100606 |
| CD4 | GK1.5 | eBioscience | 11-0041-85 |
| CD8α | 53-6.7 | TONBO biosciences | 35-0081 |
| CD25 | PC61.5 | TONBO biosciences | 65-0251 |
| CD44 | IM7 | BioLegend | 103030 |
| CD45.1 | A20 | TONBO biosciences | 75-0453 |
| CD45.2 | 104 | BioLegend | 109806 |

### 2-[N-(7-nitrobenz-2-oxa-1,3-diazol-4-yl) amino]-2-deoxy-D-glucose (2-NBDG) analysis

Mice were injected with a single dose of 375 μg of 2-NBDG (Thermo, stock: 2.5 mg/ml, injected 150 μl/mouse) in the retro-orbital venous sinus. Thymus tissues and bone marrow cells were harvested 1 hour after the injection, cells were stained as described above and were analyzed using flow cytometry. 2-NBDG fluorescence was detected in the FITC channel.

### BrdU or EdU incorporation analysis

For experiments measuring proliferation of HSCs after cyclophosphamide/GCSF administration mice were injected with a dose of 5-bromo-2′-deoxyuridine (BrdU; 1 mg/10 g body mass) and maintained on 1 mg BrdU per ml drinking water for three days. HSCs were purified as described above, fixed, stained using the BrdU APC Flow Kit (BD Biosciences) and analyzed for BrdU incorporation with flow cytometry. For experiments measuring proliferation in thymocytes, mice were injected with a dose of 5-ethynyl-2′-deoxyuridine (EdU; 1 mg/10 g body mass), the mice were euthanized 2 hours after injection, and the thymus was dissociated and stained with cell surface antibodies. Cells were then fixed in 100 μl of 1% formaldehyde on ice for 20 minutes, washed in 1x PBS + 1% BSA, and permeabilized using 0.1% saponin in 1x PBS + 1% BSA for 15 minutes on ice. Cells were stained for 20 minutes at room temperature using: 100 mM Tris pH=8.5, 1 mM CuSo4, 5 μΜ Azide-Alexa Fluor 555 (Thermo), 100 mM ascorbate. Alternatively, staining was performed using the Click-iT kit (Thermo) according to the manufacturer’s instructions. Cells were washed and analyzed with flow cytometry.

### Annexin V staining

Thymocytes were stained with fluorochrome-conjugated cell surface antibodies, washed and incubated for 20 minutes at room temperature with anti-annexin V-APC (BD Biosciences) following the manufacturer’s instructions, along with propidium iodide or DAPI to stain DNA. Cells were analyzed using flow cytometry.

### Mitochondrial mass and mitochondrial membrane potential analysis

Equal numbers of dissociated thymocytes were stained with surface markers, washed and stained at 37 °C for 30 minutes in HBSS + 2% FBS + 50 μM of verapamil and 5 μM Mitotracker Green or Mitotracker DeepRed (Thermo) to assess mitochondrial mass or mitochondrial membrane potential, respectively. Median fluorescence intensity per cell was assessed using flow cytometry.

### Bone marrow reconstitution assays

Recipient mice (CD45.1) were irradiated using an XRAD 320 X-ray irradiator (Precision X-Ray Inc.) with two doses of 540 rad (1080 rad in total) delivered at least 3h apart. Bone marrow cells were injected into the retro-orbital venous sinus of anesthetized recipients. For competitive transplants, 500,000 donor (CD45.2) and 500,000 competitor (either CD45.1 or CD45.1/CD45.2) cells were mixed and transplanted. Recipient mice were maintained on antibiotic water (Baytril 0.08 mg/ml) for 1 week pre-transplantation, and for 4 weeks post-transplantation. Peripheral blood was obtained from the tail veins of recipient mice every four weeks for at least 16 weeks after transplantation. Red blood cells were lysed with ammonium chloride potassium buffer. The remaining cells were stained with antibodies against CD45.2, CD45.1, CD45R (B220), CD11b, CD3, CD4, CD8 and Gr-1, and analyzed with flow cytometry. For experiments involving *Mx1Cre-*mediated deletion post-transplantation, poly I:C was administered to recipient mice 6 weeks after transplantation.

### Leukemia cell transplantation assays

Recipient mice were sublethally irradiated using an XRAD 320 X-ray irradiator (Precision X-Ray Inc.) with half of the lethal irradiation dose. 300,000 or 500,000 thymus cells from *Mx1Cre;Pten^Δ/Δ^;Pdha1^+^* or *Mx1Cre;Pten^Δ/Δ^;Pdha1^Δ^* mice were injected into the retro-orbital venous sinus of anesthetized recipients. Mice were bled every 2-4 weeks and analyzed for the presence of T cell leukemia in the blood using flow cytometry. Mice which appeared moribund were euthanized and their bone marrow and spleen were analyzed for the presence of T cell leukemia using flow cytometry. In moribund recipient mice which received thymus cells from *Pten^Δ/Δ^;Pdha1^Δ^* donor mice, DP cells in the bone marrow or spleen were sorted to determine the *Pdha1* deletion status using qPCR. For some recipient mice it was not possible to analyze leukemia cell *Pdha1* deletion status because mice died or were euthanized before leukemia cells could be sorted. If recipient mice appeared healthy and did not show signs of leukemia in the peripheral blood, they were analyzed at 10-15 weeks post transplantation to determine the presence or absence of leukemia in the spleen, bone marrow and thymus using flow cytometry.

### Cell isolation for rare cell number metabolomics

Isolation of cells for rare cell metabolomics was performed as previously described (Agathocleous et al., 2017). Briefly, bone marrow cells were obtained by crushing femurs, tibias, vertebrae and pelvic bones with an ice-cold mortar and pestle in a 4 °C cold room or on ice in 2 ml of Ca^2+^- and Mg^2+^-free HBSS with 0.5% bovine serum albumin. Cells were filtered through a 40 μm cell strainer into 5 ml tubes on ice. For sorting hematopoietic stem and progenitor cells, paramagnetic MicroBeads against c-Kit (Miltenyi) were added to each sample, followed 1 minute later by the fluorescent antibodies used for the isolation of the relevant cell populations. Fluorochrome-conjugated antibodies were used to enable a single staining step. For experiments in which CD45^+^ and Ter-119^+^ bone marrow cells were isolated, paramagnetic MicroBeads against CD45 and Ter-119 (Miltenyi) were used followed 1 minute later by the fluorescent antibodies used for the isolation of the relevant cell populations. Antibody staining was done for 30-60 minutes on ice in a 4 °C cold room, with occasional mixing. c-Kit^+^ cells or CD45^+^/Ter119^+^ cells were enriched using a QuadroMACS manual separator (Miltenyi) in the cold room. Cells were sorted on a FACSAria flow cytometer running with a sheath fluid of 0.5× PBS, prepared fresh using Milli-Q water (Millipore), and a 70 μm nozzle in a four-way purity sort mode. The FACSAria sheath tank was washed with Milli-Q deionized water before the experiment to eliminate contaminants.

Dedicated glassware was used to make all buffers. Each sample was in HBSS + 0.5% BSA and kept at 4 °C during sorting, which lasted about 20 minutes per sample. Equal numbers of cells from each population were directly sorted into 50 μl of cold 100% acetonitrile pre-chilled on wet ice and maintained at 4 °C during sorting. The final concentration of acetonitrile after the sort was ∼80%. No serum was used during the procedure. After sorting, each sample was kept on dry ice for the duration of the experiment, and then stored at −80 °C. Samples were vortexed for 1 minute, centrifuged at 17,000 g for 15 minutes at 4 ^0^C and the supernatant transferred into LC-MS autosampler vials. Sample processing for metabolomics was performed in a HEPA-filtered PCR hood.

### Metabolite extraction and metabolomics

For metabolomics analysis, bone marrow cells were extracted in 80:20 methanol:water (v/v) or 80:20 acetonitrile:water (v/v) kept on dry-ice for at least 2 hours and vortexed for 1 minute. Samples were centrifuged at 17,000 g for 15 minutes at 4 °C and supernatant was transferred to LC-MS autosampler vials. Thymus samples (∼10 mg) were homogenized in 1.5 ml tubes on dry ice using a sterile plastic homogenizer, kept on dry ice for at least 2 hours, then vortexed for 1 minute. Samples were then centrifuged at 17,000 g for 15 minutes at 4 °C, supernatant transferred to new 1.5 ml tubes, which were centrifuged again, and supernatant was transferred to LC-MS autosampler vials. Analysis was performed with an AB Sciex 6500+ QTrap mass spectrometer coupled to a Shimadzu LC-20A UHPLC system and with a Thermo Scientific QExactive HF-X hybrid quadrupole orbitrap high-resolution mass spectrometer (HRMS) coupled to a Vanquish UHPLC. Liquid chromatography was performed using a Millipore ZIC-pHILIC column (5 µm, 2.1 × 150 mm) with a binary solvent system of 10 mM ammonium acetate in water, pH 9.8 (solvent A) and acetonitrile (solvent B) with a constant flow rate of 0.25 ml/min. The column was equilibrated with 90% solvent B. The liquid chromatography gradient was: 0–15 minutes linear ramp from 90% B to 30% B; 15–18 minutes isocratic flow of 30% B; 18–19 minutes linear ramp from 30% B to 90% B; 19–27 column regeneration with isocratic flow of 90% B. Mass spectrometry was performed as previously described(Agathocleous et al., 2017; Tasdogan et al., 2020). For data acquired with the AB Sciex 6500, chromatogram peak areas were integrated using Multiquant (AB Sciex). For data acquired with the QExactive, chromatogram peak areas were integrated using Tracefinder (Thermo Scientific). The peak area for each metabolite was normalized to the median peak area of all the metabolites in each sample. Metabolomics data analysis, hierarchical clustering, volcano plots, and PCA plots were generated using MetaboAnalyst and Graphpad Prism.

### GSH/GSSG measurement by LC-MS/MS

For measurements of GSH and GSSG, ∼10 mg of thymus sample was placed in 80:20 methanol:water (v/v) and 0.1% formic acid on dry ice, and were manually homogenized using a plastic pestle. Samples were kept on dry ice for at least 2 hours, vortexed for 1 minute and centrifuged at 17,000 g for 15 minutes at 4 °C. The supernatant was transferred to a new tube and lyophilized using a Speedvac (Thermo Scientific). Dried metabolites were reconstituted in 0.1% formic acid in water, with 2 μl of 10 mM GSH and 2 μl of 100 μM GSSG internal standards. Reconstituted samples were vortexed, centrifuged at 17,000 g for 15 minutes at 4 °C and the supernatant was analyzed using LC-MS/MS with a polar-RP HPLC column (150 × 2 mm, 4 μm, 80 Å, Phenomenex) and the following gradient: The liquid chromatography gradient was: 0–3 minutes 0% B; 3-8 minutes linear ramp from 0% B to 100% B; 8-13 minutes isocratic flow of 100% B; return to 0% B for 5 minutes to re-equilibrate the column. Mobile Phase A was 0.1% formic acid in water. Mobile Phase B was 0.1% formic acid in acetonitrile. The flow rate was 0.2 ml/min, the column was at 4 °C and the samples in the autosampler were at 4 °C. Mass spectrometry was performed with a triple quadrupole mass spectrometer (AB Sciex QTRAP 6500).

### NAD^+^/NADH measurement by LC-MS/MS

Qualitative analysis of NAD^+^/NADH levels was performed on a QExactive HF-X mass spectrometer (Thermo Scientific, Bremen, Germany).. Quantitative analysis of NAD^+^/NADH was performed on a 6500+ mass spectrometer (AB Sciex, Framingham, MA) using a method adapted from prior work(Fu et al., 2019; Lu et al., 2018). To prepare quantitative samples, 10-20 mg thymus tissue was quickly lysed on ice using a plastic homogenizer in a tube containing 200 μl of 40:40:20 acetonitrile:methanol:water (v/v) with 0.1 M formic acid. Samples were vortexed for 10 seconds, then chilled on ice for 3 minutes. Each tube was neutralized with 17.4 μl of 15% NH_4_HCO3 (w/v) in water, then quickly vortexed and placed on dry ice for 20 minutes. Samples were centrifuged at 17,000 g for 15 minutes at 4 °C, and the supernatant was collected. ^15^N_5_-AMP internal standard was added to each extract at a final concentration of 100 μM. Samples were analyzed on a triple quadrupole mass spectrometer (AB Sciex QTRAP 6500) on the day of extraction to prevent metabolite degradation. Separation of analytes was achieved using a reversed phase C18 column (Waters HSS T3, 50 x 2.1 mm, 1.8 µm) on an Eksigent LC20A UHPLC module and a binary gradient composed of water/methanol (95:5, v/v) with 4 mM dibutylammonium acetate (DBAA) (solvent A) and water/acetonitrile (25:75, v/v, solvent B). The liquid chromatography gradient was: 0–3.2 minutes linear ramp from 0% B to 80% B; 3.2-5.2 minutes linear ramp from 80% B to 100%; 5.2-6.4 minutes isocratic flow of 100% B; return to 0% B for 1.5 minutes to re-equilibrate the column. Solvent was flowed at a constant rate of 0.15 mL/minute. The mass spectrometer was operated in multiple reaction monitoring (MRM) mode with polarity switching. Source gasses and mass transitions were optimized manually for all analytes with a T-infusion of a purified standard and mobile phase with a total flow rate of 0.15 uL/min. We optimized the following source conditions: the curtain gas was set to 20, and the collision gas was set to high. The ion spray voltage was set to −4500 in negative mode and 5500 in positive mode. The source temperature was set to 400. Ion source gas 1 was set to 20 while ion source gas 2 was set to 5. We found NAD^+^ best ionized in positive ionization mode while NADH best ionized in negative ionization mode. The ^15^N_5_-AMP internal standard was monitored in both positive and negative ionization modes. Two transitions were monitored for each analyte to confirm its identity but only one transition was used for quantitation. For NAD^+^, 664/428 and 664/524 were monitored; 664/428 was used for quantitation. For NADH, 664/397 and 664/408 were monitored; 664/408 was used for quantitation. For ^15^N_5_-AMP, 353/141 was monitored in positive mode while 351/139 was monitored in negative mode. For quantitation, NAD^+^ signal was normalized to the ^15^N_5_-AMP signal in positive mode, while the NADH signal was normalized to the ^15^N_5_-AMP signal in negative mode. All cellular extracts were analyzed against an 8-point standard curve ranging from 5 nM to 1000 nM. All standard curves had R^2^ values greater than or equal to 0.98 with greater than 6 calibrators having accuracies within 20% of their known concentration.

### Pyruvate and α-ketoglutarate measurement by LC-MS/MS

For pyruvate and α-ketoglutarate measurement by derivatization as indicated in the figures, ∼ 10 mg of thymus or pelleted bone marrow cells were lysed in 50 μl of ice cold buffer containing 50:50 (v/v) methanol:water, 100 μM DETAPAC-HCl, 1% formic acid and 0.3% NaN_3_. After three rounds of freeze/thaw cycles in liquid nitrogen, samples were sonicated on ice for 1 minute. Samples were incubated for 1 hour at −20 °C, vortexed and protein was pelleted via centrifugation at 20,000 g at 4 °C for 15 minutes. 40 μl of the supernatant was derivatized with 4 μl of 500 μM o-phenylenediamine or 4-methoxy-o-phenylenediamine (final concentration 50 μM) for 4 hours at room temperature protected from light in 1.5 ml tubes. Then, samples were centrifuged at 20,000 g for 15 minutes, and supernatants were transferred to glass vials for LC-MS/MS analysis. The mass spectrometer was an AB Sciex 6500+ operating in multiple reaction monitoring (MRM) mode. Liquid chromatography was performed using a Scherzo SM-C18 column (Imtakt). Mobile phase A was 0.1% formic acid in water and mobile phase B 0.1% formic acid in acetonitrile. The flow rate was 0.3 ml/min. The liquid chromatography gradient was: 0–2 minutes 0% B; 2-6 minutes linear ramp from 0% B to 100% B; 6-8 minutes isocratic flow of 100% B; 8-8.5 minutes linear ramp to 0% B. 8.5-11 minutes 0% B. A valve switch was used to allow flow into the mass spectrometer between 3-7.5 minutes. Pyruvate was monitored in the positive mode with the transitions 133/92 and 133/65 (o-phenylenediamine derivatization) or 163/122 and 163/95 (4-methoxy-o-phenylenediamine derivatization) and α-ketoglutarate with the transitions 191/173 and 191/131 (o-phenylenediamine derivatization) or 221/203 and 221/161 (4-methoxy-o-phenylenediamine derivatization).

### *Ex vivo* U^13^C-glucose tracing

1∼2 million DP cells were sorted into a 10 ml tube containing 5 ml of HBSS + 2% FBS and were kept on ice until the end of sorting. Cells were washed with 1x PBS at 1500 rpm for 5 minutes at 4 °C, were resuspended in 1 ml of the tracing media before plating them on a 12 well plate. For bone marrow cells, each femur was flushed with 3 ml of HBSS + 2% FBS, 5∼8 million cells were washed with 1x PBS at 1500 rpm for 5 minutes at 4 °C, and were resuspended with 1 ml of the tracing media before plating them on a 12 well plate. Tracing media was composed of: DMEM (Sigma, D5030) with 18 mM glucose (U^13^C-glucose labeled glucose for the experiment and unlabeled glucose for untraced control), 4 mM glutamine, and 10% dialyzed FBS (Hyclone). Untraced control samples were composed of a mix of dissociated thymocytes and flushed bone marrow cells (1:1). The samples were incubated in tracing media for 2 hours at 37 °C. At the 2 hour time point, cells were collected and centrifuged at 1500 rpm for 5 minutes at 4 °C, washed with 1X ice-cold saline, extracted in 100μl of 40:40:20 acetonitrile:methanol:water (v/v) and stored in −80 °C. On the day of extraction, samples were vortexed for 1 minute, underwent three rounds of freeze-thaw cycles to extract the metabolites, followed by centrifugation at 16,000 g for 15 minutes at 4 °C. The supernatant was collected and analyzed on a triple quadrupole mass spectrometer (AB Sciex QTRAP 6500) using LC-MS/MS on the day of extraction. LC-MS analysis was performed using HILIC chromatography as described in the metabolomics section. The mass spectrometer operated in MRM mode. Transitions used were: acetyl-CoA (positive mode): M+0 810/303, M+1 811/304, M+2 812/305; citrate (negative mode): M+0 191/111, M+1 192/111 and 192/112, M+2 193/112 and 193/113, M+3 194/113 and 194/114, M+4 195/114 and 195/115, M+5 196/115 and 196/116; M+6 197/116.

### *In vivo* U^13^C-glucose tracing

Mice were fasted for 16 hours before the start of infusions. Mice were placed under anesthesia using 40 μl injection of 25 mg/ml ketamine/xylazine and a 27-gauge catheter was inserted in the lateral tail vein. Mice were infused with U^13^C-glucose (CLM-1396, Cambridge Isotope Laboratories), starting with a bolus of 0.4125 mg g^−1^ body mass over 1 minute in 125 μl of saline, followed by continuous infusion of 0.008 mg g^−1^ body mass per minute for 3 hours (in a volume of 150 μl.h^−1^)(Faubert et al., 2017). Plasma samples were collected at 0, 30, 60, 120, and 180 minutes of infusion. 3 hours after the start of infusion, mice were sacrificed, thymus tissues collected and immediately snap frozen in liquid nitrogen. For bone marrow, each femur was immediately flushed with 200 μl of ice-cold 80:20 methanol:water (v/v) and snap frozen in liquid nitrogen.

### Rare cell *in vivo* U^13^C-glucose tracing using LC-MS/MS

U^13^C-glucose tracing in mice was performed as described in another section. To determine changes in labeling during cell purification, we compared thymus samples which were snap frozen, or bone marrow cells which were extracted by flushing femurs with ice-cold 80:20 methanol:water immediately after cessation of tracing, *versus* purified thymocytes or CD45^+^ or Ter119^+^ bone marrow cells. Cell purification was performed as described in the section for rare cell metabolomics with the exception that 1x PBS was used as a cell buffer in place of 1x HBSS. LC-MS/MS analysis was performed using HILIC chromatography as described in the metabolomics section, coupled to an AB Sciex 6500+ triple quadrupole mass spectrometer operating in MRM mode. Labeled transitions were determined by analyzing metabolites from cells cultured with U^13^C-glucose for 24 hours. Transitions used were: Positive mode: acetyl-carnitine: M+0 204/85, M+1 205/85, M+2 206/85; carnitine: M+0 162/103, M+1 163/103, M+2 164/103; aspartate: M+0 134/74, M+1 135/74 and 135/75, M+2 136/74, 136/75 and 136/76, M+3 137/75 and 137/76, M+4 138/76; serine: M+0 106/60, M+1 107/60 and 107/61, M+2 108/61 and 108/62, M+3 109/62; glutamine: M+0 147/84, M+1 148/84 and 148/85, M+2 149/85 and 149/86, M+3 150/86 and 150/87, M+4 151/87, 151/88 and 152/88; glycerophosphocholine M+0 258/104, M+1 259/104, M+2 260/104 and M+3 261/104. Negative mode: Glutamate M+0 146/102, M+1 147/102 and 147/103, M+2 148/103 and 148/104, M+3 149/104 and 149/105, M+4 150/105 and 150/106, M+5 151/106. Data analysis was performed with MultiQuant (AB Sciex). For each experiment, bone marrow or thymus from an untraced mouse was analyzed in parallel as a control. Typically, in stable isotope tracing experiments mass isotopologue distributions are corrected for natural abundance. However in this method, which had a mass resolution of 1, we found that untraced samples sometimes contained co-eluting peaks from compounds that were detected in the same transition as an isotopomer of a targeted molecule, which complicated use of natural abundance correction algorithms. An additional consideration was that, because the method operated at the limit of sensitivity, the linearity assumed by correction algorithms may not hold at the lowest signal intensities. For those two reasons, we preferred to correct for the presence of naturally abundant isotopes by subtracting signal in the untraced sample from that of the traced samples. To calculate the glucose-derived labeling for each metabolite, the fractional abundance for each isotopologue was normalized to the fractional abundance of U^13^C-glucose in the plasma.

### Gas chromatography mass spectrometry (GC-MS/MS)

Thymus tissues weighing 10-30 mg were homogenized using an electronic tissue disruptor (Qiagen) in ice-cold 80:20 methanol:water (v/v) followed by three freeze-thaw cycles in liquid nitrogen. Samples were centrifuged at 13,000 g at 4°C for 15 minutes, and supernatant was collected and lyophilized. Bone marrow samples were vortexed for 1 minute, underwent 3 freeze-thaw cycles in liquid nitrogen, followed by centrifugation at 13,000 g at 4 °C for 15 minutes. Supernatant was collected and lyophilized. For plasma sample preparation, whole blood was chilled on ice during collection then centrifuged at 13,000 g at 4 °C for 15 minutes to separate the plasma. 5 μl of plasma were added to ice-cold 80:20 methanol:water (v/v), and were lyophilized. All thymus, bone marrow and plasma samples were lyophilized using a SpeedVac (Thermo) and re-suspended in 20 μl of methoxyamine (10 mg/ml) in pyridine made fresh every time. Samples were then incubated at 70 °C for 15 minutes and centrifuged at 500 g at 4 °C for 5 minutes. Supernatant was transferred to autoinjector vials containing 40 μl of N-(tert-butyldimethylsilyl)-N-methyltrifluoroacetamide (MTBSTFA) to derivatize polar metabolites. Samples were incubated for 1 hour at 70 °C, then aliquots of 2 μl were injected for analysis using Agilent 7890 gas chromatograph coupled to a 5975C Mass Selective Detector. The observed distributions of mass isotopologues were corrected for natural abundance as previously described(Yang et al., 2014). To calculate the glucose-derived labeling for each metabolite, the fractional abundance for each isotopologue was normalized to the fractional abundance of U^13^C-glucose in the plasma. Each data point represents the average of two technical replicates from each mouse.

### Protein extraction and western blot analysis

Equal numbers of cells from each population were sorted into tubes containing HBSS+2%FBS media and then washed with ice-cold 1X PBS. Pelleted cells were extracted either in 1% Triton or RIPA buffer containing phosphatase and protease inhibitors. Samples were vortexed every 10 minutes on ice for 30 minutes and centrifuged at 16,000 g for 10 minutes. PLB loading buffer (BioRad) and BME were added to the supernatant and samples were heated at 70 °C for 10 minutes with agitation. Samples were separated on 4-15% Mini-Protean TGX gels (BioRad) and transferred to 0.2 μm PVDF membranes or nitrocellulose membranes (BioRad) by wet transfer using Tris Glycine transfer buffer (BioRad). Western blots were performed using antibodies against PDH (Abcam, ab110330), vinculin (CST, 4650S), H3K27ac (CST, 8173S), H3 (CST, 9715S), H4K16ac (Abcam, ab109463), and H3K9ac (CST, 9649S). In cases where the loading control antigen was the same size as the antigen we probed for (for example in histone acetylation experiments) we ran parallel gels from the same samples for loading controls to avoid stripping and re-probing. For low numbers of cells (less than 30,000 cells), the SuperSignal Western Blot Enhancer kit (Thermo Scientific) was used according to the manufacturer’s instructions. Signals were detected using the SuperSignal West Pico or SuperSignal West Femto chemiluminescence kits (Thermo Scientific). Western blot signals were quantified using ImageJ and protein signal of interest was normalized to the respective loading control.

### RNA extraction and quantitative real-time PCR (qRT-PCR) analysis of bone marrow

Total RNA was isolated from cells using Trizol (Invitrogen). Complementary DNA (cDNA) was synthesized with 1-5 μg of RNA using iScript cDNA synthesis kit (Bio-Rad). mRNA levels were measured using qRT-PCR with SYBR Green Master Mix (BioRad). mRNA levels were normalized to β-actin expression. To account for the possibility of genomic DNA (gDNA) contamination, no reverse-transcriptase (no-RT) control reaction was used. cDNA extracted from testis was used as a positive control for *Pdha2* expression.

#### Primers used for RT-PCR or qPCR experiments

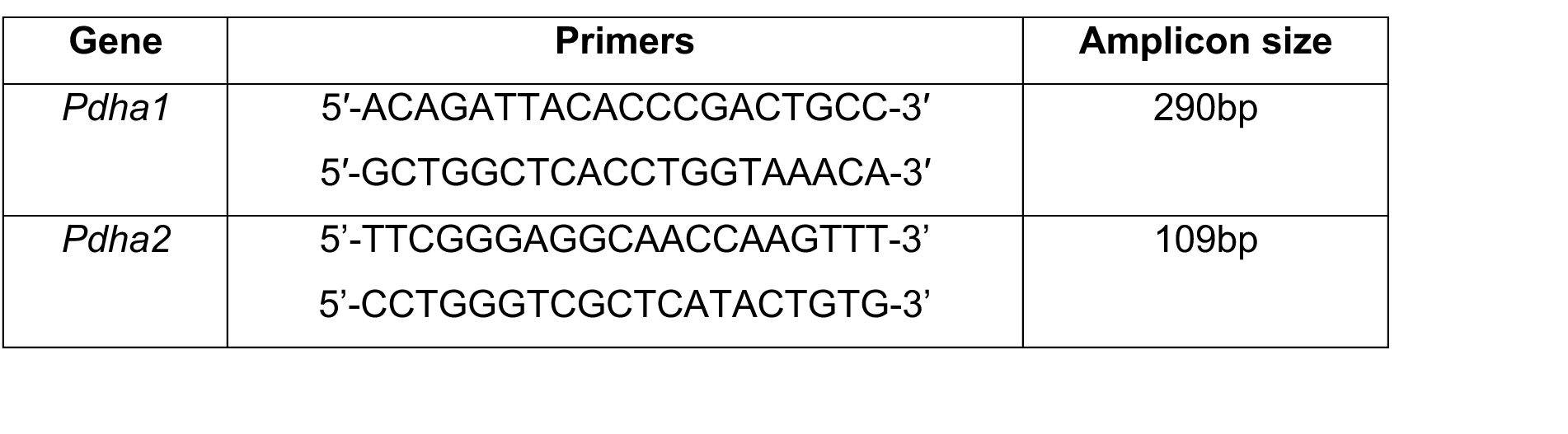
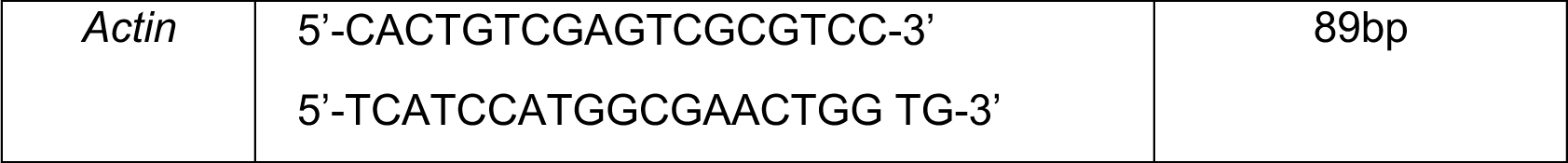

#### Primers used for PCR reactions

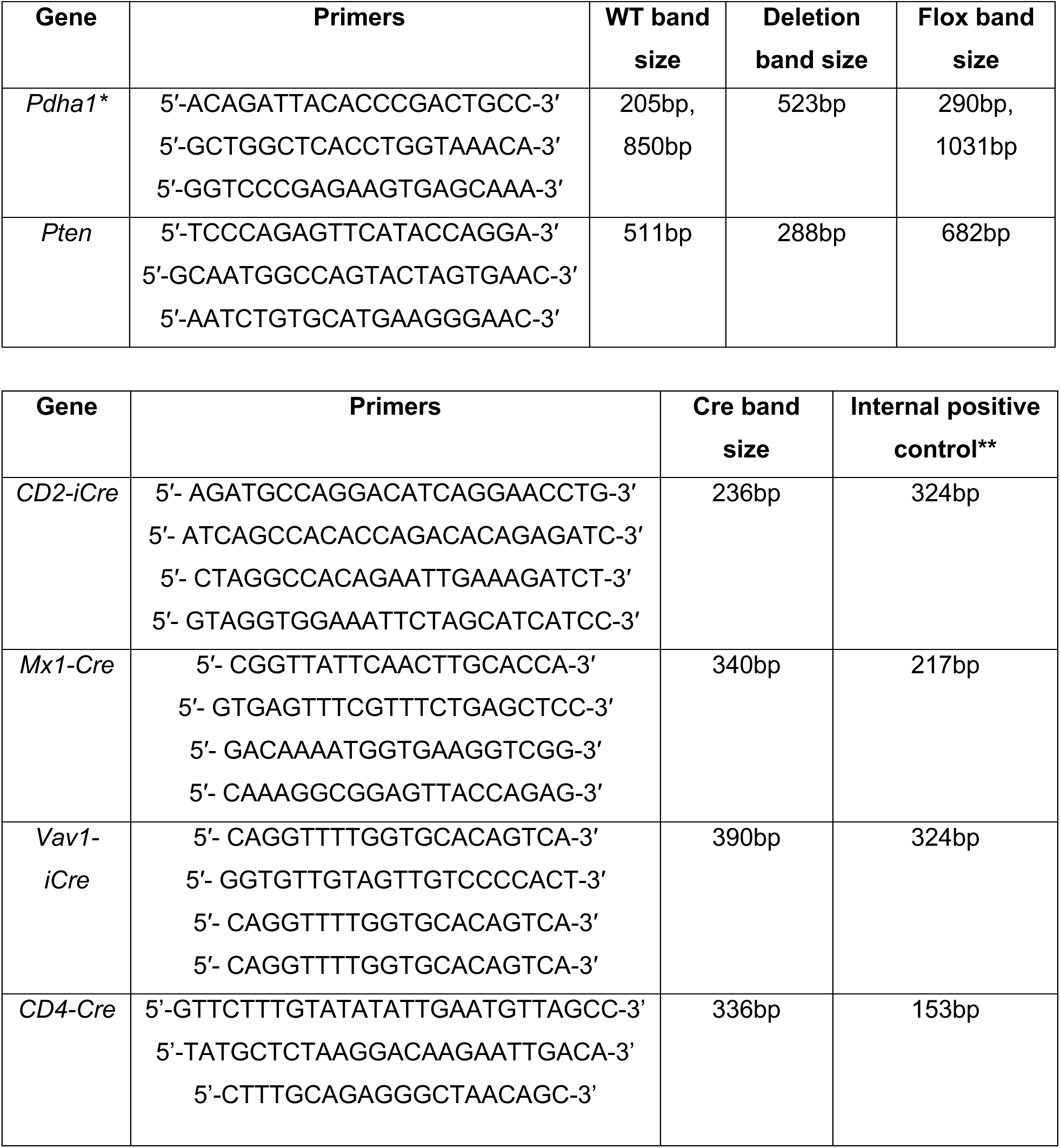

### Statistical Analysis

Each figure panel shows the pooled results from all the mice we analyzed from multiple independent experiments. Mice were allocated to experiments randomly. For most experiments the operator was not blinded to the genotypes. Littermate controls, or controls from litters of the same parental strains born a few days apart were used for experiments. Prior to analyzing the statistical significance of differences among treatments, we tested whether the data were normally distributed and whether variance was similar among treatments. To test for normal distribution, we performed the Shapiro–Wilk test when 3≤n<20 or the D’Agostino Omnibus test when n≥20. To test if variability significantly differed among treatments, we performed F-tests (for experiments with two treatments) or Levene’s median tests (for more than two treatments). If the data did not significantly deviate from normality, we used a parametric test, otherwise data were log-transformed and tested for a significant deviation from normality. If the log-transformed data did not significantly deviate from normality, a parametric test was used on the transformed data. If both the untransformed data and log-transformed data significantly deviated from normality a non-parametric test was used on the untransformed data. To assess the statistical significance of a difference between two treatments, we used a t-test for data which did not significantly deviate from normality and did not have significantly unequal variances, or a t-test with Welch’s correction for data which did not significantly deviate from normality but had significantly unequal variances, or a Mann-Whitney U test for unpaired or Wilcoxon matched-pairs test for paired data which significantly deviated from normality. To assess the statistical significance of a difference between more than two treatments, we used a one-way ANOVA for data which did not significantly deviate from normality and did not have significantly unequal variances, a Brown-Forsythe ANOVA for data which did not significantly deviate from normality and had significantly unequal variances, and a Kruskal-Wallis test for data which significantly deviated from normality. To assess the statistical significance of differences between treatments when multiple measurements across time were taken for each treatment, we used a two-way ANOVA (e.g. for the proportion of a hematopoietic cell type in different genotypes analyzed using different mice at various time points after deletion), or a two-way ANOVA with repeated measures and the Geisser-Greenhouse correction (e.g. for the case of transplantation experiments where blood from the same mouse was sampled repeatedly over time), or a repeated-measures mixed-effects model (when datapoints were missing). To assess differences in survival between cohorts of mice, we used a logrank Mantel-Cox test. We corrected for multiple comparisons by controlling the false discovery rate at 5% using the method of Benjamini, Krieger and Yekutieli (in the case of multiple t-tests) or by using the Holm-Sidak test (for ANOVA tests where variance was not significantly different between samples) or Dunnett T3 test (for ANOVA tests where variance was significantly different between samples) or Dunn’s test (for non-parametric tests). All statistical analyses were performed with Graphpad Prism 8.3.

---

**Supplementary Figure 1.**
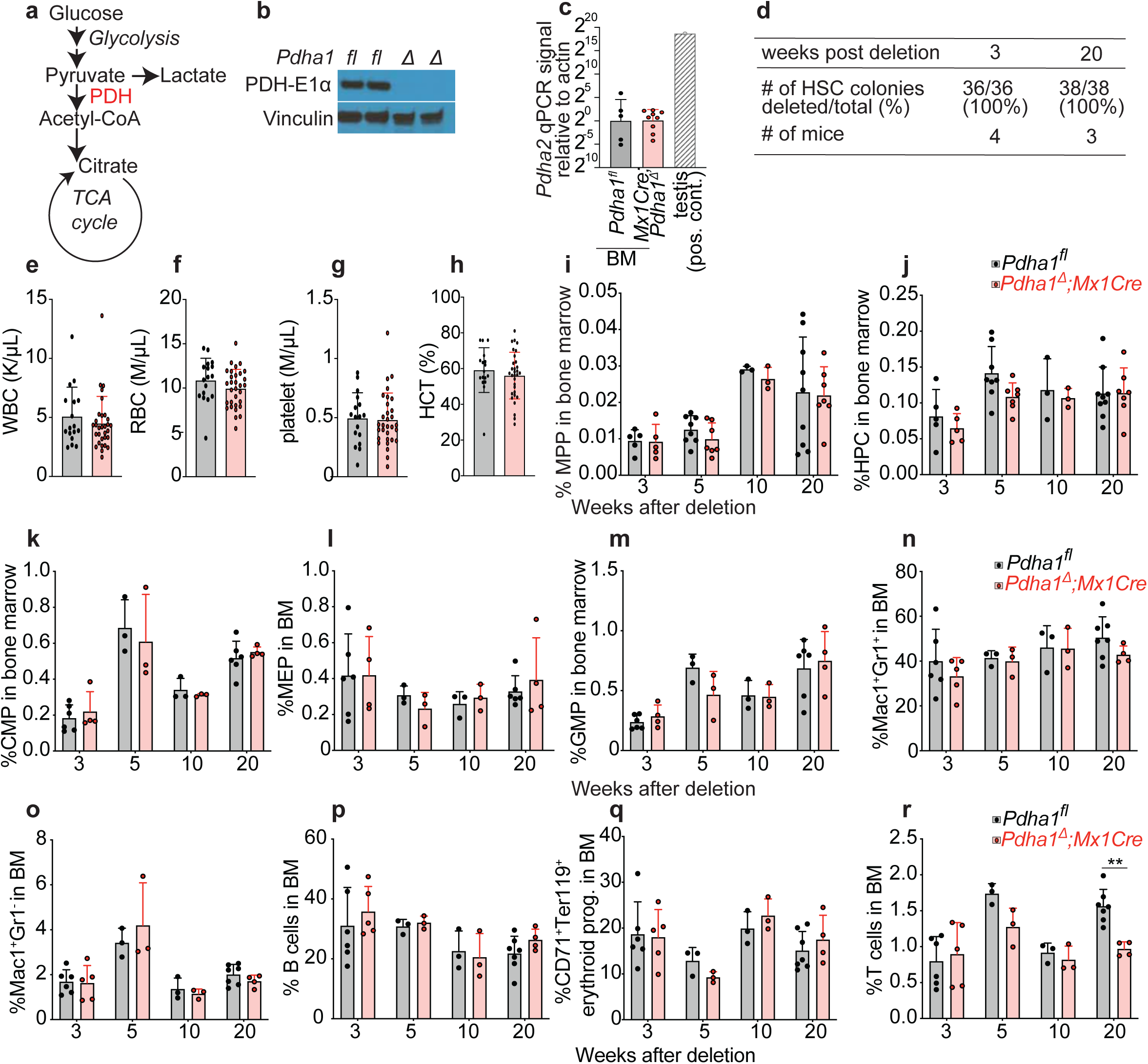
Hematopoietic analysis in the bone marrow of *Mx1Cre*;*Pdha1^Δ^* mice. **(a)** Simplified diagram of glycolysis and TCA cycle, showing their connection through PDH. **(b)** Western blot against PDH-E1, which is encoded by *Pdha1,* was performed using bone marrow cells from *Mx1Cre*;*Pdha1^Δ^* mice and littermate controls (representative blot shown from a total of n=7-8 mice per genotype from 3 independent experiments). **(c)** RT-qPCR of Pdha2, a testis-specific homologue of Pdha1, in the bone marrow of *Mx1Cre*;*Pdha1^Δ^* mice and littermate controls (n=5-9 per genotype, n=1 for testis positive control). The qPCR signal for Pdha2 was >2^15^ lower for bone marrow cells as compared to testis, as measured by cycle difference from actin amplification. The number of cycles required to amplify Pdha2 in the bone marrow in RT-qPCR was not lower than the number of cycles in the no-RT controls, suggesting the low signal in the bone marrow originates from genomic DNA rather than Pdha2 transcripts. **(d)** Genotyping of colonies grown in culture from single HSCs purified from *Mx1Cre*;*Pdha1^Δ^* mice showing that 100% of HSCs have been deleted. **(e-h)** Analysis of the blood of *Pdha1^fl^* and *Pdha1^Δ^* mice (n=17-31 mice per genotype). **(i-r)** Bone marrow analysis of *Mx1Cre*;*Pdha1^Δ^* mice and littermate controls (n=3-9 mice/genotype/time-point). All data represent mean ± s.d. *p < 0.05, **p < 0.01, ***p < 0.001. Statistical significance in (r) was assessed with a two-way ANOVA and in (c) with a one-way ANOVA.

**Supplementary Figure 2.**
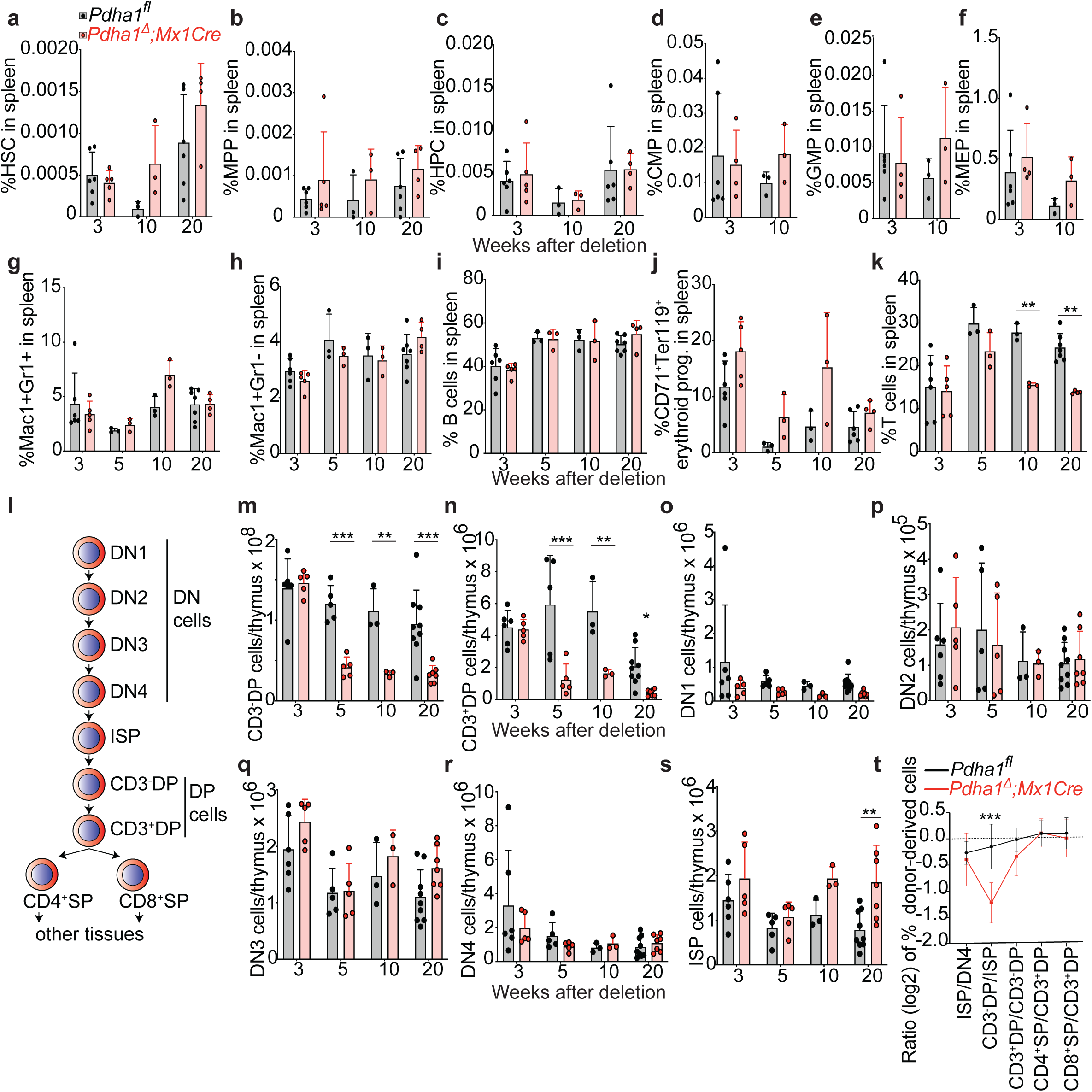
Hematopoietic analysis in the spleen and thymus of *Mx1Cre*;*Pdha1^Δ^* mice. **(a-k)** Analysis of the spleen of *Mx1Cre*;*Pdha1^Δ^* mice and littermate controls (n=3-6 mice/genotype/time point). **(l)** Overview of T cell lineage development in the thymus. CD3^-^DP cells are the most abundant of the T cell progenitors (> 90%). **(m-s)** Analysis of the thymus of *Mx1Cre*;*Pdha1^Δ^* mice and littermate controls (n=3-9 mice/genotype/time-point). **(t)** Results from competitive transplantation experiments of *Mx1Cre;Pdha1^Δ^* and littermate control mice along with wild type competitors into lethally irradiated recipients. Depicted is the proportion of donor-derived cells for each stage of thymocyte development divided by the proportion of donor-derived cells from the preceding stage (n=6-11 mice per genotype). All data represent mean ± s.d. *p < 0.05, **p < 0.01, ***p < 0.001. Statistical significance was assessed with a two-way ANOVA (k, m, n, s), or multiple t-tests (p).

**Supplementary Figure 3.**
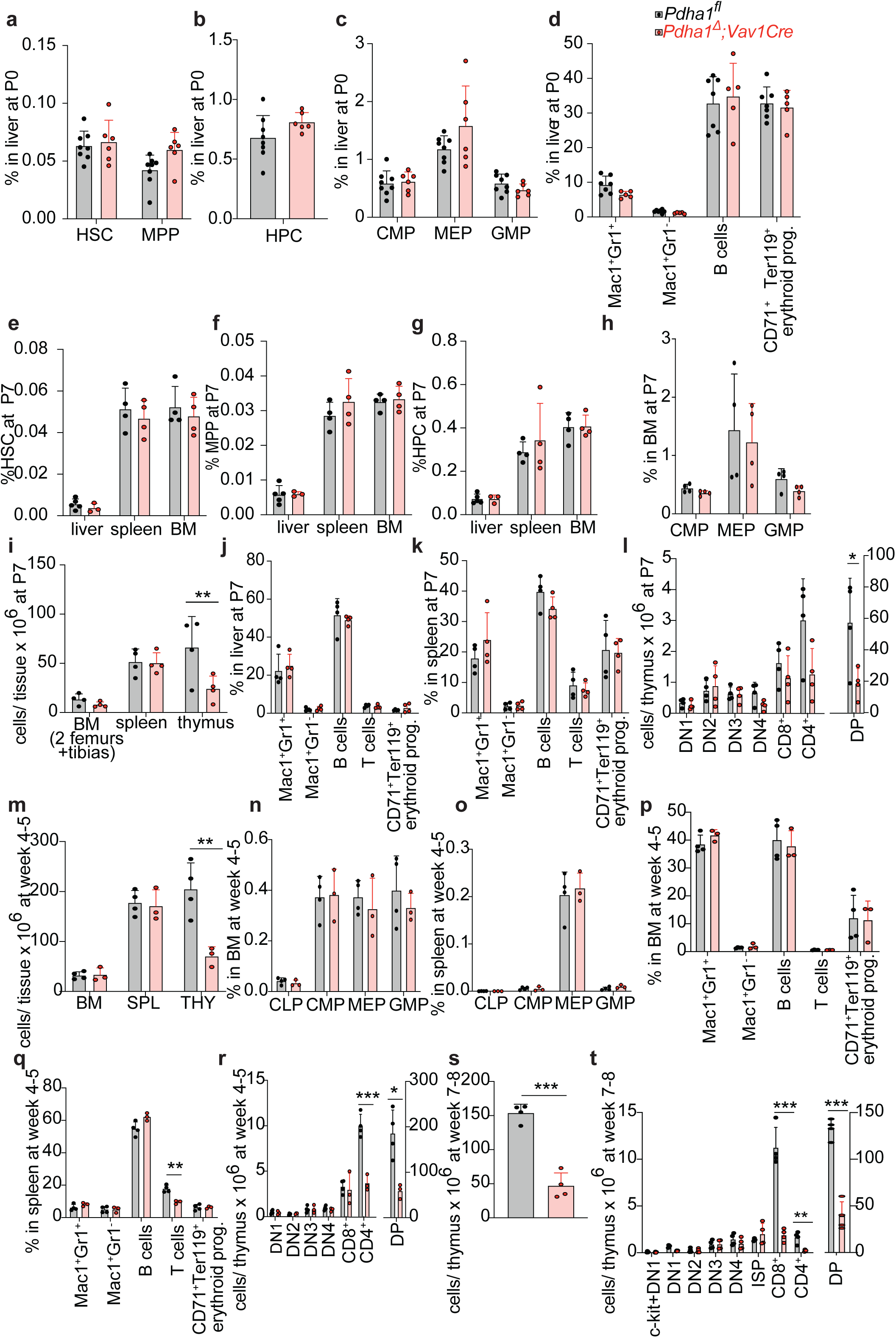
Hematopoietic analysis in the liver, spleen, bone marrow and thymus of *Vav1Cre*;*Pdha1^Δ^* mice at different postnatal ages. Analysis of hematopoiesis in the liver, spleen, bone marrow and thymus of *Vav1Cre*;*Pdha1^Δ^* mice and littermate controls at P0 (**a-d**, n=6-8 mice per genotype), P7 (**e-l**, n=4 mice per genotype) and week 4-5 of age (**m-r**, n=3-4 mice per genotype). **(s-t)** Analysis of T lymphopoiesis in the thymus of *Vav1Cre*;*Pdha1^Δ^* mice and littermate controls at week 7-8 of age (n=4 mice per genotype). All data represent mean ± s.d. *p < 0.05, **p < 0.01, ***p < 0.001. Statistical significance was assessed with a t-test (i, l, m, q, r, s, t-CD4^+^ cells), t test with Welch’s correction (t-CD8^+^ cells), or Mann-Whitney (t-DP cells).

**Supplementary Figure 4.**
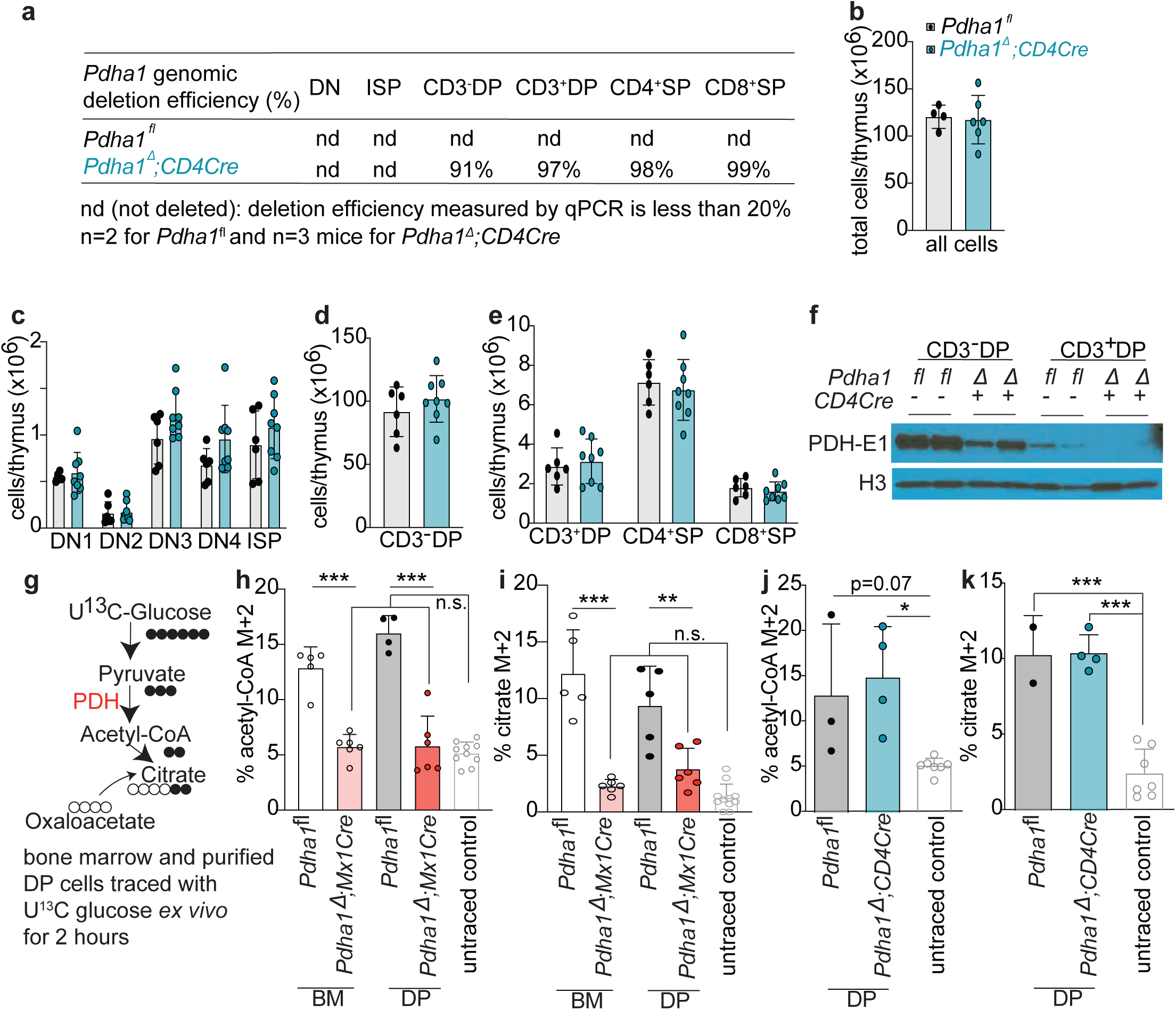
The effect of *CD4Cre*-mediated *Pdha1* deletion on T cell lymphopoiesis and metabolism. **(a)** Percentage of *Pdha1* deleted alleles as assessed by qPCR analysis of purified thymus cell types in *CD4Cre*;*Pdha1^Δ^* mice and littermate controls (n=2-3 mice per genotype). **(b-e)** Analysis of T lymphopoiesis in the thymus of *CD4Cre*;*Pdha1^Δ^* mice and littermate controls (n=6-8 mice per genotype). **(f)** Western blots against PDH-E1 using purified CD3^-^DP and CD3^+^DP cells from *CD4Cre*;*Pdha1^Δ^* mice or littermate controls (shown is a representative blot from a total of n=5-6 mice/genotype from 2 independent experiments). **(g)** Schematic of *ex vivo* tracing using U^13^C-glucose. Bone marrow and purified DP cells were traced for 2 hours at 37^0^C, and labeling was measured by LC-MS/MS. **(h-k)** Acetyl-CoA (M+2) and citrate (M+2) labeling in bone marrow cells and in purified DP cells from *Mx1Cre*;*Pdha1^Δ^* mice and *CD4Cre*;*Pdha1^Δ^* mice (n=3-10 mice/genotype or untraced control). All data represent mean ± s.d. *p < 0.05, **p < 0.01, ***p < 0.001. Statistical significance was assessed using t-tests (h, i) and one-way ANOVA (j, k). t-tests or one-way ANOVA was performed on log-transformed values if they were normally distributed.

**Supplementary Figure 5.**
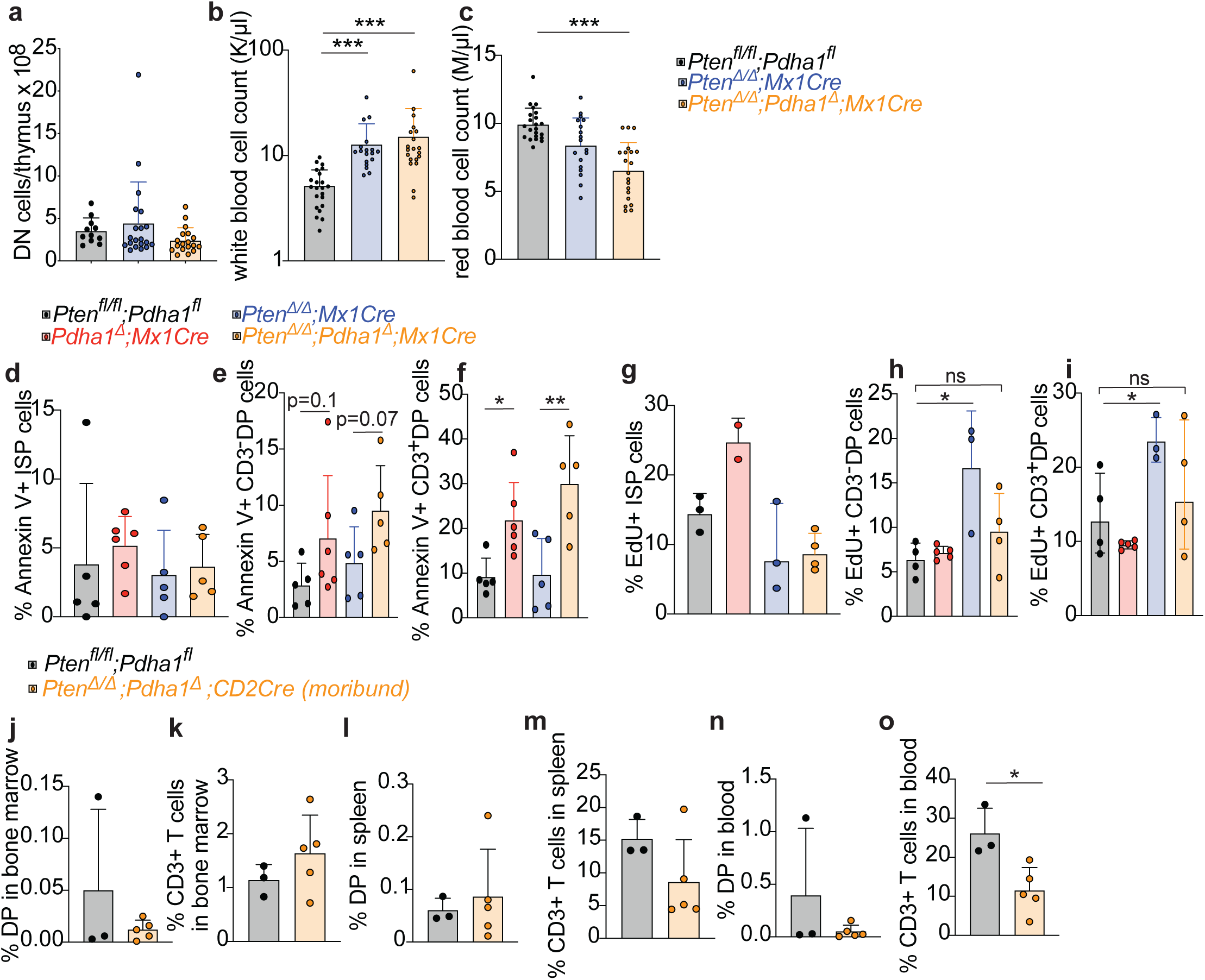
Hematopoietic analysis in *Mx1Cre*;*Pten^Δ/Δ^*;*Pdha1^Δ^* and *Mx1Cre*; *Pdha1^Δ^* mice, and analysis of moribund *CD2Cre*;*Pten ^Δ/Δ^*;*Pdha1^Δ^* mice. **(a)** Number of DN thymocytes in *Mx1Cre*;*Pten^Δ/Δ^*;*Pdha1^Δ^* mice and littermate controls (n= 11-20 mice/genotype). **(b-c)** Analysis of blood cells in *Mx1Cre*;*Pten^Δ/Δ^*;*Pdha1^Δ^* mice and littermate controls (n=18-22 mice/genotype). **(d-f)** % of each cell type that was positive for Annexin V staining using flow cytometry in thymus of wild-type, *Mx1Cre*;*Pdha1^Δ^*, *Mx1Cre*;*Pten ^Δ/Δ^* and *Mx1Cre*;*Pten ^Δ/Δ^*;*Pdha1^Δ^* mice 4-6 weeks after poly(I:C) administration (n=4-6 mice per genotype). **(g-i)** % of each cell type that was positive for 5-ethynyl-2-deoxyuridine (EdU) incorporation measured by flow cytometry in thymus of wild-type, *Mx1Cre*;*Pdha1^Δ^*, *Mx1Cre*;*Pten^Δ/Δ^* and *Mx1Cre*;*Pten^Δ/Δ^*;*Pdha1^Δ^* mice (n=2-4 mice per genotype). **(j-o)** Moribund *CD2Cre*;*Pten^Δ/Δ^*;*Pdha1^Δ^* mice analyzed at 10-21 months of age did not have T cell leukemia, indicated by normal or decreased number of T cells in the bone marrow, spleen, and blood, and an absence of disseminated DP cells (n=5 mice moribund *CD2Cre*;*Pten^Δ/Δ^*;*Pdha1^Δ^* mice and 3 wild type controls). All data represent mean ± s.d. *p < 0.05, **p < 0.01, ***p < 0.001. Statistical significance was assessed using Kruskal-Wallis test (b, c), t-tests (e, f, h, o), and t-tests with Welch correction (i).

**Supplementary Figure 6.**
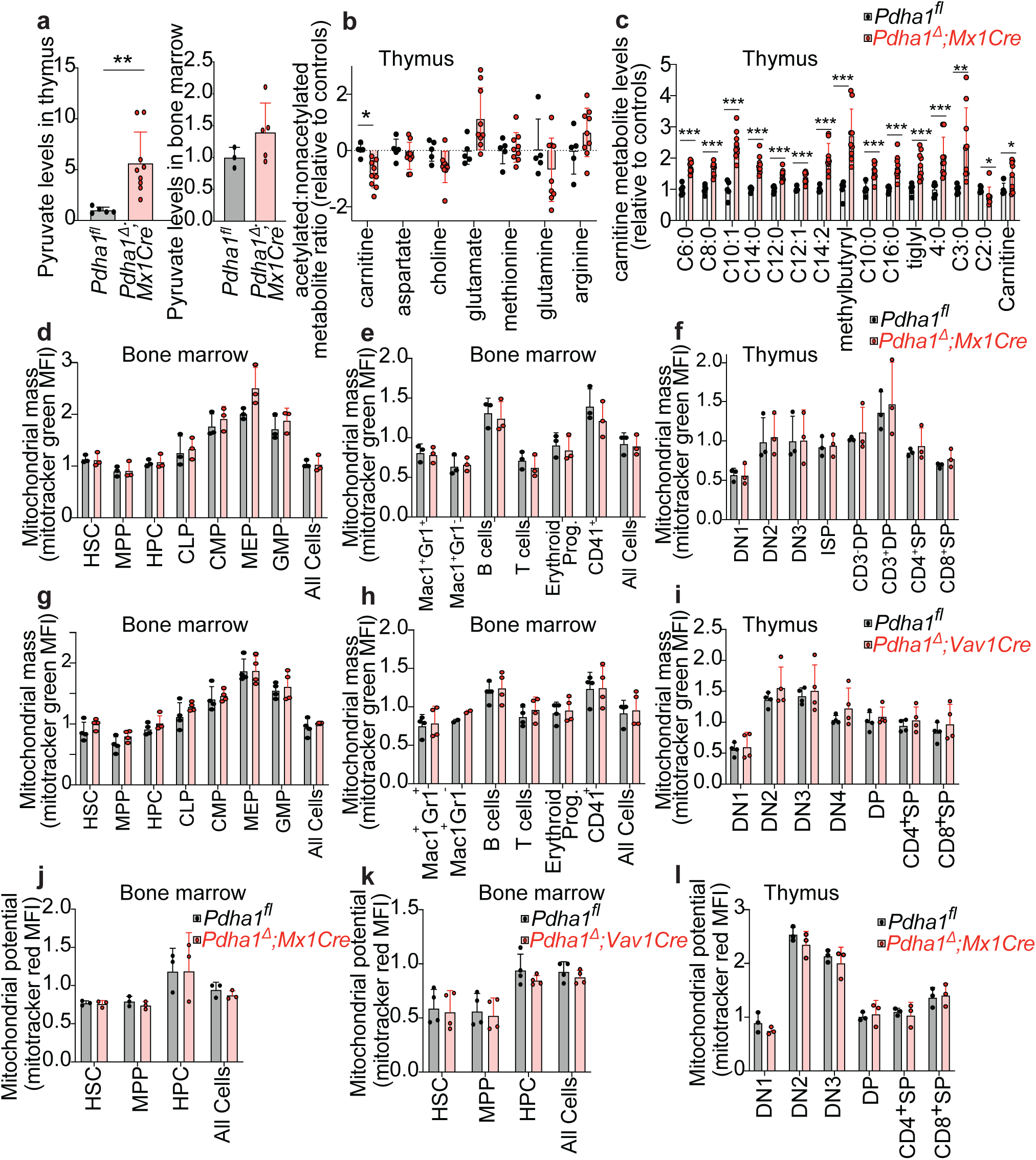
Hematopoietic *Pdha1* deletion elevates pyruvate in the thymus but not in bone marrow, and does not change acetylated metabolites, and mitochondrial mass and potential. **(a)** Pyruvate levels in the thymus and bone marrow of *Mx1Cre*;*Pdha1^Δ^* mice and littermate controls (n=3-9 mice/genotype/tissue). Measurements are derived from metabolomics experiments. **(b)** Ratio of acetylated:non-acetylated metabolites for each metabolite noted in the x-axis in the thymus of *Mx1Cre*;*Pdha1^Δ^* mice and littermate controls, relative to the value for the controls (n=5-9 mice/genotype). **(c)** Levels of carnitines in the thymus of *Mx1Cre*;*Pdha1^Δ^* mice and littermate controls, relative to the value for the controls (n=5-9 mice/genotype). **(d-i)** Mitochondrial mass of different hematopoietic cell types in the bone marrow and thymus of *Mx1Cre*;*Pdha1^Δ^* (n=3) or *Vav1Cre*;*Pdha1^Δ^* (n=4) mice and littermate controls (n=3-4), as measured by median fluorescence intensity of Mitotracker Green. **(j-l)** Mitochondrial membrane potential of different hematopoietic cell types in the bone marrow and thymus of *Mx1Cre*;*Pdha1^Δ^* (n=3), *Vav1Cre*;*Pdha1^Δ^* mice (n=4) and littermate controls (n=3-4), as measured by median fluorescence intensity of Mitotracker Red. All data represent mean ± s.d. *p < 0.05, **p < 0.01, ***p < 0.001. Statistical significance was assessed using t-tests (a, b, c). Multiple comparisons correction was performed by controlling the false discovery rate at 5%.

**Supplementary Figure 7.**
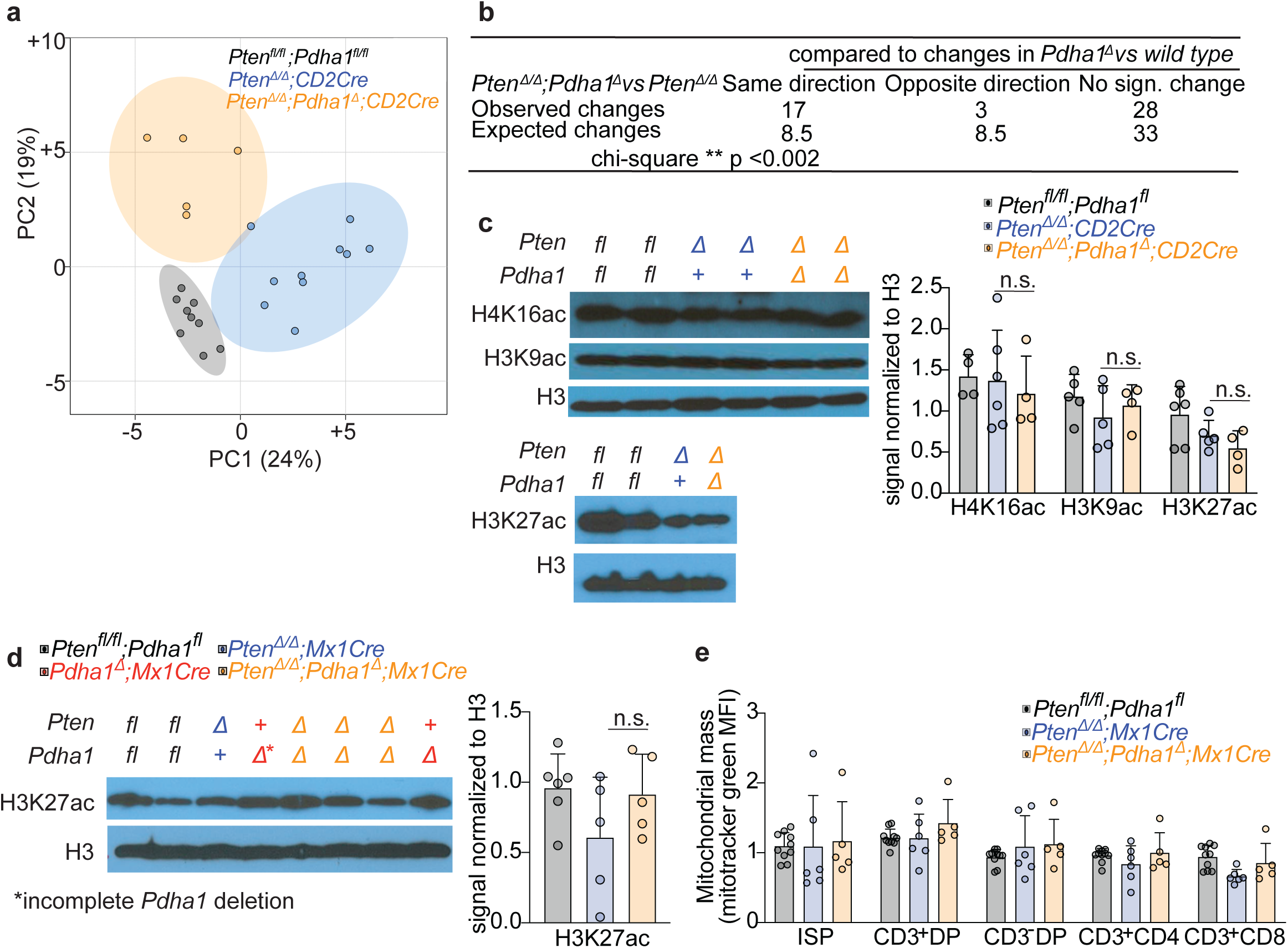
Metabolic and histone acetylation changes caused by PDH deficiency in the context of *Pten* deletion. **(a)** Unbiased clustering using a PCA plot of metabolomic data from the thymus of mice from the indicated genotypes (n=5-10 mice/genotype) showing that the thymus of each genotype is metabolically distinct. Each dot represents a sample from a different mouse. **(b)** Table summarizing the overlap between the metabolic effects of PDH-deficiency in wild type and *Pten-* deficient thymus. A total of 50 metabolites significantly changed in *CD2Cre;Pten^Δ/Δ^*;*Pdha1^Δ^* as compared to *CD2Cre;Pten^Δ/Δ^* thymus. Of those, we observed that 17 changed in the same direction in *Mx1Cre;Pdha1^Δ^* as compared to *Pdha1^fl^* (wild type) thymus, 3 in the opposite direction, 28 did not change and 2 were not detected. The number of observed changes that changed in the same direction between *Pten^Δ/Δ^*;*Pdha1^Δ^ vs Pten^Δ/Δ^* and *Pdha1^Δ^ vs* wild type (17) was significantly higher than would be expected by chance, as determined by taking into account that 34% of metabolites significantly changed in *Pdha1^Δ^ vs* wild type thymus. **(c-d)** Western blots against H4K16ac, H3K9ac, and H3K27ac in purified DP cells from *CD2Cre*;*Pten ^Δ/Δ^*;*Pdha1^Δ^* (c) or *Mx1Cre;Pten^Δ/Δ^*;*Pdha1^Δ^* (d) mice and littermate controls (a total of n=4-6 mice/genotype. Representative blots from 2-3 independent experiments are shown). **(e)** Mitochondrial mass measurement using mitotracker green in thymocytes from *Mx1Cre*;*Pten ^Δ/Δ^*;*Pdha1^Δ^* mice and littermate controls (n=5-10 mice per genotype). All data represent mean ± s.d. *p < 0.05, **p < 0.01, ***p < 0.001. Statistical significance was assessed using a chi-square test (b) and two-way ANOVA (c,d).

**Supplementary table 1.** Metabolic changes in the thymus after *Pdha1* deletion

|  | fold change<br>*Pdha1*<sup>Δ</sup> / *Pdha1*<sup>f/l</sup> | p value | q value |
| --- | --- | --- | --- |
| Cystathionine | 0.13 | <0.0001 | 0.0003 |
| Carnitine (6:0) | 1.73 | <0.0001 | 0.0008 |
| Quinolate | 0.40 | <0.0001 | 0.0016 |
| Carnitine (8:0) | 1.66 | <0.0001 | 0.0016 |
| Carnitine (10:1) | 2.42 | <0.0001 | 0.0016 |
| Deoxycytidine | 0.39 | 0.0001 | 0.0018 |
| GSH | 0.66 | 0.0001 | 0.0021 |
| Carnitine (14:0) | 1.79 | 0.0001 | 0.0021 |
| Glutamate | 0.46 | 0.0002 | 0.0039 |
| N-Acetyl-D-galactosamine 4-sulfate | 0.49 | 0.0002 | 0.0039 |
| N-Acetylglutamine | 0.31 | 0.0002 | 0.0044 |
| N-Arachidonoyl dopamine | 0.27 | 0.0003 | 0.0047 |
| Carnitine (12:0) | 1.45 | 0.0003 | 0.0047 |
| Hypotaurine | 0.49 | 0.0003 | 0.0052 |
| Carnitine (12:1) | 1.39 | 0.0004 | 0.0053 |
| Carnitine (14:2) | 2.01 | 0.0005 | 0.0069 |
| UDP | 0.71 | 0.0006 | 0.0069 |
| ADP | 0.61 | 0.0007 | 0.0077 |
| Carnitine (methylbutyryl) | 2.71 | 0.0011 | 0.0121 |
| 5-Methylcytosine | 0.61 | 0.0014 | 0.0140 |
| 2-hydroxyglutarate | 1.83 | 0.0015 | 0.0140 |
| Carnitine (10:0) | 1.57 | 0.0015 | 0.0140 |
| Hexosamine phosphate | 0.61 | 0.0016 | 0.0140 |
| dATP | 0.32 | 0.0016 | 0.0140 |
| Carnitine (16:0) | 1.70 | 0.0017 | 0.0140 |
| 2'-O-Methylinosine | 0.59 | 0.0018 | 0.0147 |
| Acetoin | 4.92 | 0.0019 | 0.0149 |
| 2-Aminoadipate | 0.38 | 0.0020 | 0.0152 |
| N-Ribosylnicotinamide | 0.48 | 0.0024 | 0.0172 |
| Carnitine (tiglyl) | 1.78 | 0.0025 | 0.0179 |
| Deoxyuridine | 0.64 | 0.0030 | 0.0207 |
| 1-Pyrroline-5-carboxylate | 2.15 | 0.0035 | 0.0229 |
| GSSG | 0.23 | 0.0036 | 0.0235 |
| Carnitine (4:0) | 2.03 | 0.0045 | 0.0283 |
| N-Glycolylneuraminate | 1.44 | 0.0047 | 0.0287 |
| UDP-Hexose | 0.67 | 0.0057 | 0.0334 |
| Proline | 1.49 | 0.0063 | 0.0354 |
| Pyruvate | 5.62 | 0.0063 | 0.0354 |
| N-Acetylaspartylglutamate | 0.38 | 0.0066 | 0.0359 |
| 5-hydroxy-2,4-dioxopentanoate | 0.69 | 0.0075 | 0.0397 |
| Saccharate | 0.66 | 0.0091 | 0.0463 |
| CMP-N-glycolylneuraminate | 0.67 | 0.0092 | 0.0463 |
| GDP | 0.59 | 0.0111 | 0.0547 |
| Glycerol 3-phosphate | 0.63 | 0.0129 | 0.0619 |
| Aconitate | 0.79 | 0.0131 | 0.0619 |
| Methylnicotinic acid | 0.71 | 0.0146 | 0.0673 |
| Phosphoserine | 5.76 | 0.0150 | 0.0675 |
| UDP-N-acetylhexosamine | 0.75 | 0.0157 | 0.0688 |
| epsilon-(gamma-Glutamyl)-lysine | 1.49 | 0.0162 | 0.0688 |
| Threonate | 0.46 | 0.0162 | 0.0688 |
| Glutamine | 0.54 | 0.0177 | 0.0715 |
| Ascorbic acid 2-sulfate | 0.49 | 0.0180 | 0.0715 |
| Dimethyllysine | 0.66 | 0.0183 | 0.0715 |
| Dihydroorotate | 0.44 | 0.0185 | 0.0715 |
| N-Acetylornithine | 1.99 | 0.0185 | 0.0715 |
| Propionylcarnitine | 2.44 | 0.0189 | 0.0717 |
| Choline | 1.47 | 0.0203 | 0.0745 |
| Deoxyadenosine | 0.31 | 0.0204 | 0.0745 |
| dCMP | 0.58 | 0.0226 | 0.0814 |
| Orotidine | 0.71 | 0.0249 | 0.0882 |
| Glycerophosphoglycerol | 0.71 | 0.0257 | 0.0889 |
| Lactate | 1.39 | 0.0260 | 0.0889 |
| Hexose-phosphate | 1.86 | 0.0269 | 0.0903 |
| NADP+ | 0.68 | 0.0272 | 0.0903 |
| Carboxyethylarginine | 30.91 | 0.0281 | 0.0916 |
| Carnosine | 1.82 | 0.0286 | 0.0918 |
| Sedoheptulose 7-phosphate | 5.89 | 0.0293 | 0.0926 |
| Fructose lysine-6-phosphate | 0.70 | 0.0327 | 0.1019 |
| Allantoin | 0.78 | 0.0350 | 0.1077 |
| Isoleucine | 0.78 | 0.0357 | 0.1083 |
| N-Acetyl-4-O-acetylneuraminate | 0.37 | 0.0377 | 0.1120 |
| Glucosylgalactosyl hydroxylysine | 0.45 | 0.0386 | 0.1120 |
| Ribitol phosphate | 3.45 | 0.0386 | 0.1120 |
| S-Methyl-thioadenosine | 0.64 | 0.0391 | 0.1120 |
| Sedoheptulose | 0.64 | 0.0439 | 0.1206 |
| 2-Hydroxy-3-oxopropanoate | 3.56 | 0.0441 | 0.1206 |
| Creatinine | 0.81 | 0.0442 | 0.1206 |
| Lysine | 0.64 | 0.0444 | 0.1206 |
| Glycylglutamine | 1.51 | 0.0461 | 0.1237 |
| Carnitine (6:1) | 1.38 | 0.0467 | 0.1238 |
| dCTP | 0.43 | 0.0494 | 0.1281 |
| Levulinate | 1.77 | 0.0495 | 0.1281 |

**Supplementary table 2.**
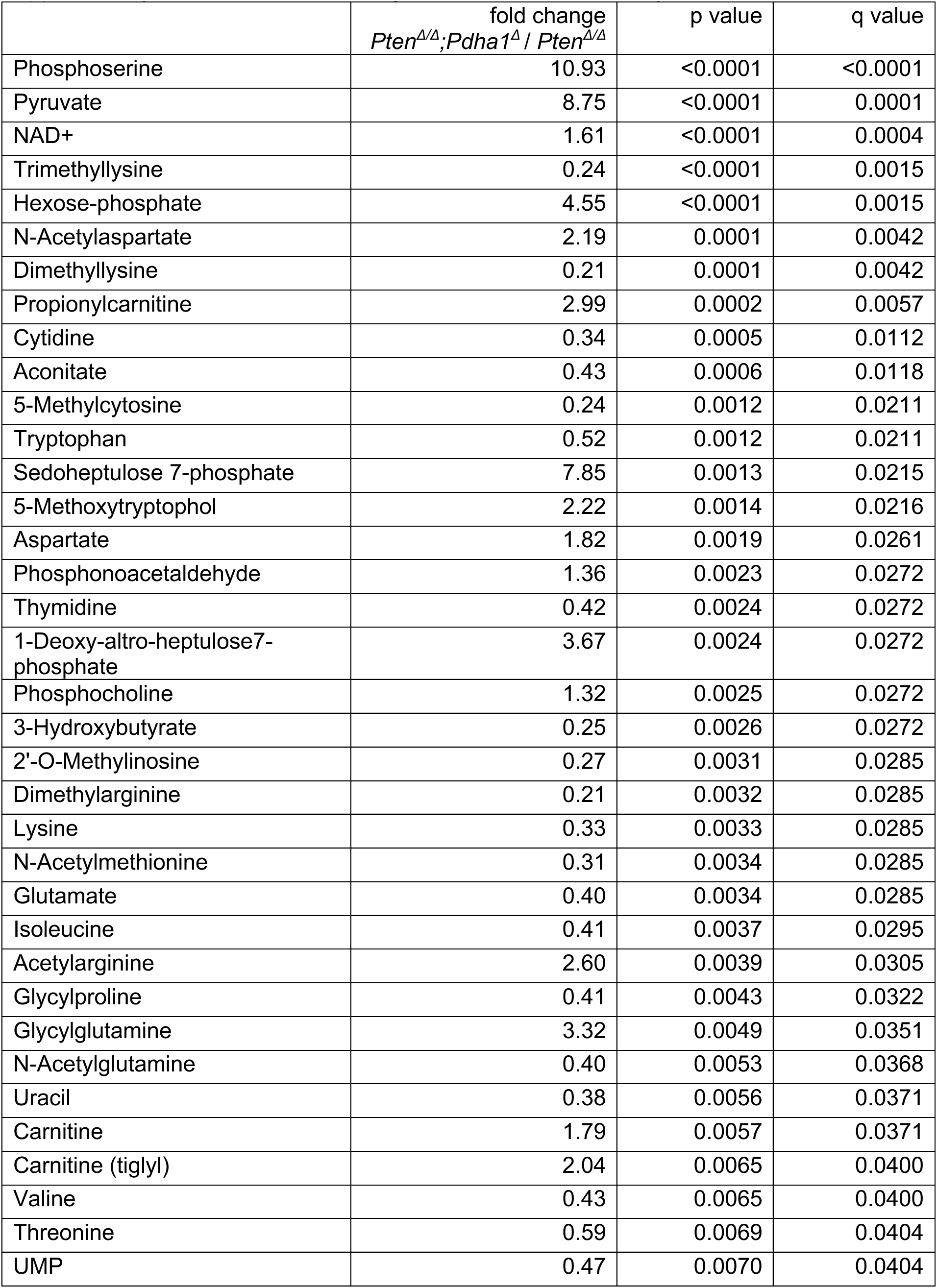
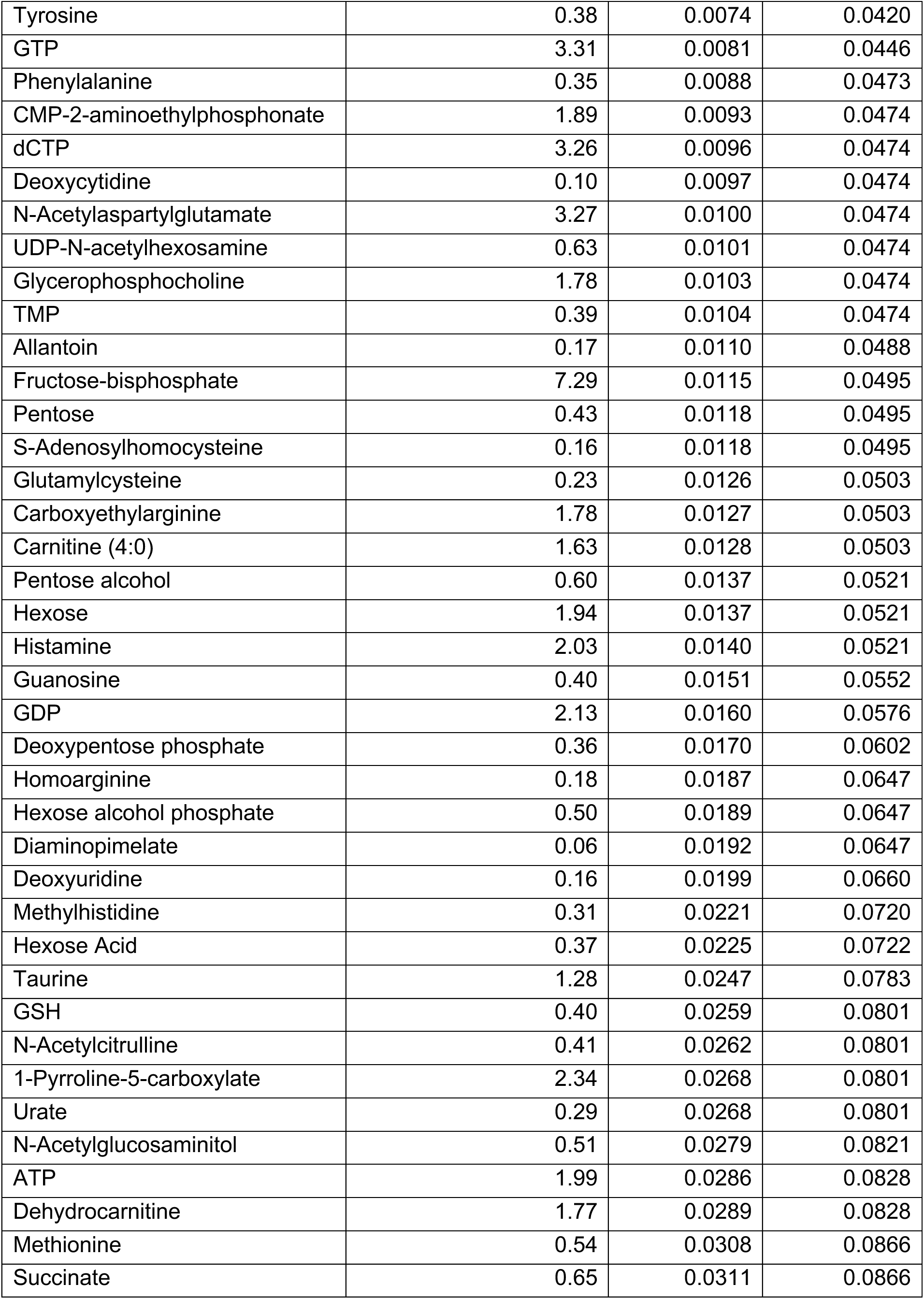
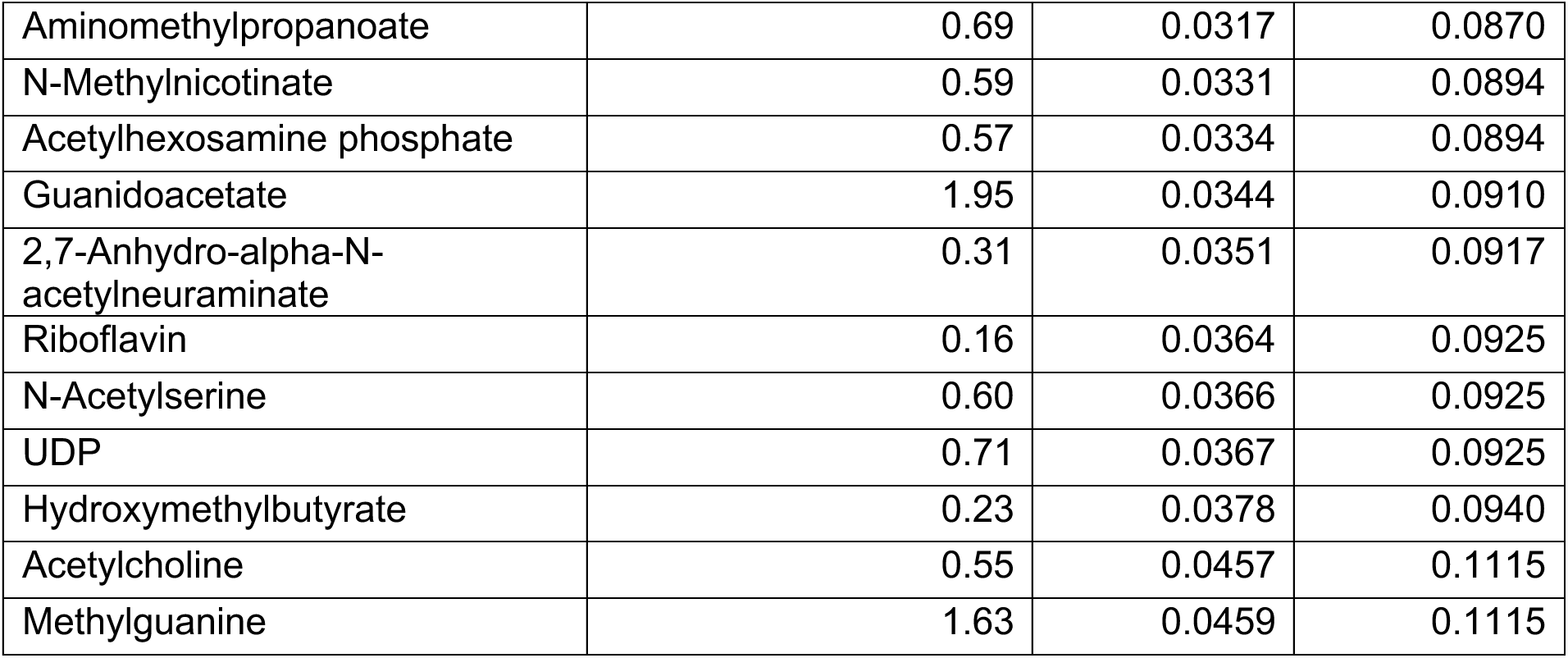
Metabolic changes in the *Pten-*deficient thymus after *Pdha1* deletion

|  | fold change<br>*Pten*<sup>Δ/Δ</sup>; *Pdha1*<sup>Δ</sup> / *Pten*<sup>Δ/Δ</sup> | p value | q value |
| --- | --- | --- | --- |
| Phosphoserine | 10.93 | <0.0001 | <0.0001 |
| Pyruvate | 8.75 | <0.0001 | 0.0001 |
| NAD+ | 1.61 | <0.0001 | 0.0004 |
| Trimethyllysine | 0.24 | <0.0001 | 0.0015 |
| Hexose-phosphate | 4.55 | <0.0001 | 0.0015 |
| N-Acetylaspartate | 2.19 | 0.0001 | 0.0042 |
| Dimethyllysine | 0.21 | 0.0001 | 0.0042 |
| Propionylcarnitine | 2.99 | 0.0002 | 0.0057 |
| Cytidine | 0.34 | 0.0005 | 0.0112 |
| Aconitate | 0.43 | 0.0006 | 0.0118 |
| 5-Methylcytosine | 0.24 | 0.0012 | 0.0211 |
| Tryptophan | 0.52 | 0.0012 | 0.0211 |
| Sedoheptulose 7-phosphate | 7.85 | 0.0013 | 0.0215 |
| 5-Methoxytryptophol | 2.22 | 0.0014 | 0.0216 |
| Aspartate | 1.82 | 0.0019 | 0.0261 |
| Phosphonoacetaldehyde | 1.36 | 0.0023 | 0.0272 |
| Thymidine | 0.42 | 0.0024 | 0.0272 |
| 1-Deoxy-altro-heptulose7-phosphate | 3.67 | 0.0024 | 0.0272 |
| Phosphocholine | 1.32 | 0.0025 | 0.0272 |
| 3-Hydroxybutyrate | 0.25 | 0.0026 | 0.0272 |
| 2'-O-Methylinosine | 0.27 | 0.0031 | 0.0285 |
| Dimethylarginine | 0.21 | 0.0032 | 0.0285 |
| Lysine | 0.33 | 0.0033 | 0.0285 |
| N-Acetylmethionine | 0.31 | 0.0034 | 0.0285 |
| Glutamate | 0.40 | 0.0034 | 0.0285 |
| Isoleucine | 0.41 | 0.0037 | 0.0295 |
| Acetylarginine | 2.60 | 0.0039 | 0.0305 |
| Glycylproline | 0.41 | 0.0043 | 0.0322 |
| Glycylglutamine | 3.32 | 0.0049 | 0.0351 |
| N-Acetylglutamine | 0.40 | 0.0053 | 0.0368 |
| Uracil | 0.38 | 0.0056 | 0.0371 |
| Carnitine | 1.79 | 0.0057 | 0.0371 |
| Carnitine (tiglyl) | 2.04 | 0.0065 | 0.0400 |
| Valine | 0.43 | 0.0065 | 0.0400 |
| Threonine | 0.59 | 0.0069 | 0.0404 |
| UMP | 0.47 | 0.0070 | 0.0404 |
| Tyrosine | 0.38 | 0.0074 | 0.0420 |
| GTP | 3.31 | 0.0081 | 0.0446 |
| Phenylalanine | 0.35 | 0.0088 | 0.0473 |
| CMP-2-aminoethylphosphonate | 1.89 | 0.0093 | 0.0474 |
| dCTP | 3.26 | 0.0096 | 0.0474 |
| Deoxycytidine | 0.10 | 0.0097 | 0.0474 |
| N-Acetylaspartylglutamate | 3.27 | 0.0100 | 0.0474 |
| UDP-N-acetylhexosamine | 0.63 | 0.0101 | 0.0474 |
| Glycerophosphocholine | 1.78 | 0.0103 | 0.0474 |
| TMP | 0.39 | 0.0104 | 0.0474 |
| Allantoin | 0.17 | 0.0110 | 0.0488 |
| Fructose-bisphosphate | 7.29 | 0.0115 | 0.0495 |
| Pentose | 0.43 | 0.0118 | 0.0495 |
| S-Adenosylhomocysteine | 0.16 | 0.0118 | 0.0495 |
| Glutamylcysteine | 0.23 | 0.0126 | 0.0503 |
| Carboxyethylarginine | 1.78 | 0.0127 | 0.0503 |
| Carnitine (4:0) | 1.63 | 0.0128 | 0.0503 |
| Pentose alcohol | 0.60 | 0.0137 | 0.0521 |
| Hexose | 1.94 | 0.0137 | 0.0521 |
| Histamine | 2.03 | 0.0140 | 0.0521 |
| Guanosine | 0.40 | 0.0151 | 0.0552 |
| GDP | 2.13 | 0.0160 | 0.0576 |
| Deoxypentose phosphate | 0.36 | 0.0170 | 0.0602 |
| Homoarginine | 0.18 | 0.0187 | 0.0647 |
| Hexose alcohol phosphate | 0.50 | 0.0189 | 0.0647 |
| Diaminopimelate | 0.06 | 0.0192 | 0.0647 |
| Deoxyuridine | 0.16 | 0.0199 | 0.0660 |
| Methylhistidine | 0.31 | 0.0221 | 0.0720 |
| Hexose Acid | 0.37 | 0.0225 | 0.0722 |
| Taurine | 1.28 | 0.0247 | 0.0783 |
| GSH | 0.40 | 0.0259 | 0.0801 |
| N-Acetylcitrulline | 0.41 | 0.0262 | 0.0801 |
| 1-Pyrroline-5-carboxylate | 2.34 | 0.0268 | 0.0801 |
| Urate | 0.29 | 0.0268 | 0.0801 |
| N-Acetylglucosaminitol | 0.51 | 0.0279 | 0.0821 |
| ATP | 1.99 | 0.0286 | 0.0828 |
| Dehydrocarnitine | 1.77 | 0.0289 | 0.0828 |
| Methionine | 0.54 | 0.0308 | 0.0866 |
| Succinate | 0.65 | 0.0311 | 0.0866 |
| Aminomethylpropanoate | 0.69 | 0.0317 | 0.0870 |
| N-Methylnicotinate | 0.59 | 0.0331 | 0.0894 |
| Acetylhexosamine phosphate | 0.57 | 0.0334 | 0.0894 |
| Guanidoacetate | 1.95 | 0.0344 | 0.0910 |
| 2,7-Anhydro-alpha-N-acetylneuraminate | 0.31 | 0.0351 | 0.0917 |
| Riboflavin | 0.16 | 0.0364 | 0.0925 |
| N-Acetylserine | 0.60 | 0.0366 | 0.0925 |
| UDP | 0.71 | 0.0367 | 0.0925 |
| Hydroxymethylbutyrate | 0.23 | 0.0378 | 0.0940 |
| Acetylcholine | 0.55 | 0.0457 | 0.1115 |
| Methylguanine | 1.63 | 0.0459 | 0.1115 |

